# RAB-35 regulates distinct steps of trogocytosis in the biting and bitten cell

**DOI:** 10.64898/2026.02.20.707090

**Authors:** Julie Manikas, Liam Popovsky, Yusuff Abdu, Jeremy Nance

**Affiliations:** Department of Cell Biology, NYU Grossman School of Medicine, New York, NY 10016; Center for Quantitative Cell Imaging, U. of Wisconsin, Madison, WI 53706; Department of Cell and Regenerative Biology, U. of Wisconsin, Madison, WI 53706

**Keywords:** <u>Key words</u>: trogocytosis, phagocytosis, cannibalism, germ line, scission

## Abstract

Trogocytosis is a form of cellular cannibalism in which a cell “bites” off pieces of another cell. Here, we investigate the molecular mechanisms of a developmentally programmed *C. elegans* trogocytic event that occurs when endodermal cells bite off and digest pieces of primordial germ cells (PGCs) called lobes. Through a genetic screen, we identify the Rab family small GTPase *rab-35* as a central regulator of trogocytosis and show that its function is required in both biting (endodermal) and bitten (PGC) cells. Within endodermal cells, RAB-35 enriches around trogocytosed PGC lobes, promotes the removal of phosphatidylinositol 4,5-bisphosphate (PIP_2_), and is required for lobe digestion. By contrast, we show that RAB-35 within PGCs works with the ESCRT complex to promote scission of the PGC lobe from the cell body. Our findings identify a new regulator of trogocytosis that has distinct functions in the biting and bitten cells and provide evidence that the bitten cell contributes to the scission of its own membrane.

## INTRODUCTION

Cells bite off pieces of other cells through a cannibalistic process called trogocytosis (Greek, ‘cell nibbling’). Trogocytosis was first described as the mechanism the parasitic amoeba *Naegleria fowleri* uses to kill mammalian cells (Brown, 1979). More recently, trogocytosis has been observed in many important non-pathogenic cell interactions (reviewed in (Bettadapur et al., 2020; Rodriguez et al., 2025; Zhao et al., 2022)). For example, mouse retinal pigment epithelial cells bite off and digest spent photoreceptor outer segments to prevent retinal degeneration (Umapathy et al., 2023). During development, zebrafish macrophages nibble pieces of hematopoietic stem cells to assess their fitness and trigger the removal of low-quality stem cells (Wattrus et al., 2022). In the immune system, trogocytosis can either promote or suppress an immune response (reviewed in (Guha and Banerjee, 2025; Yang et al., 2025)). For example, neutrophils clear *Trichomonias vaginalis* infections via trogocytosis (Mercer et al., 2018), and donor hematopoietic stem cells (HSCs) can trogocytose host antigens to evade immune detection (Chow et al., 2013; Yamanaka et al., 2009). Cells can also change their behavior by gaining factors through trogocytosis, such as when HSCs trogocytose the guidance receptor CXCR4 from macrophages to promote retention in the bone marrow niche (Gao et al., 2024). Finally, trogocytosis is increasingly recognized for its importance in cancer biology and treatment. For example, neutrophils can use trogocytosis to kill antibody-opsonized tumor cells (Matlung et al., 2018), and conversely, tumor cells can escape death in chimeric antigen receptor T-cell therapy (CAR-T) by using trogocytosis to transfer target antigens to other cells (Hamieh et al., 2019). Despite the importance of trogocytosis in a wide array of essential biological events, the molecular mechanisms required to execute it are poorly understood.

Trogocytosis is conceptually similar to phagocytosis, in which an engulfing cell consumes a target cell or a previously shed cellular piece in its entirety (reviewed in (Depierre et al., 2025; Lafuente et al., 2020)). However, trogocytosis differs in that an additional biting step is required to sever the cell fragment from the target cell membrane. Trogocytosis can be subdivided conceptually into three steps: target <u>recognition</u>; <u>scission</u> and internalization of the target cell fragment; and <u>digestion</u> of the internalized fragment (Fig. 1A-C). Target recognition is thought to be mediated by “eat-me” signals (reviewed in (Cockram et al., 2021; Park and Kim, 2017)) (Fig. 1A). Once a target is recognized, the biting cell uses Rac-induced F-actin-rich extensions to surround the target cell fragment (Martinez-Martin et al., 2011; Ralston et al., 2014; Umapathy et al., 2023). Next, the biting cell must cut off the bitten cell fragment (‘scission’) and reseal its own plasma membrane to fully internalize the cargo (Fig. 1B). Scission requires proteins that bind to and induce membrane curvature, such as BAR-domain proteins, to recruit the membrane-severing GTPase dynamin to the site of membrane cutting (Fig. 1B) (Abdu et al., 2016; Umapathy et al., 2023). It is unclear whether dynamin alone is sufficient to complete scission, or whether other mechanisms contribute. While there is no scission step needed in phagocytosis, dynamin is required for the engulfing cell to reseal its membrane after phagocytosis is complete – a process called phagosome sealing (Fig. 1B) (Cheng et al., 2015).

**Figure 1.**
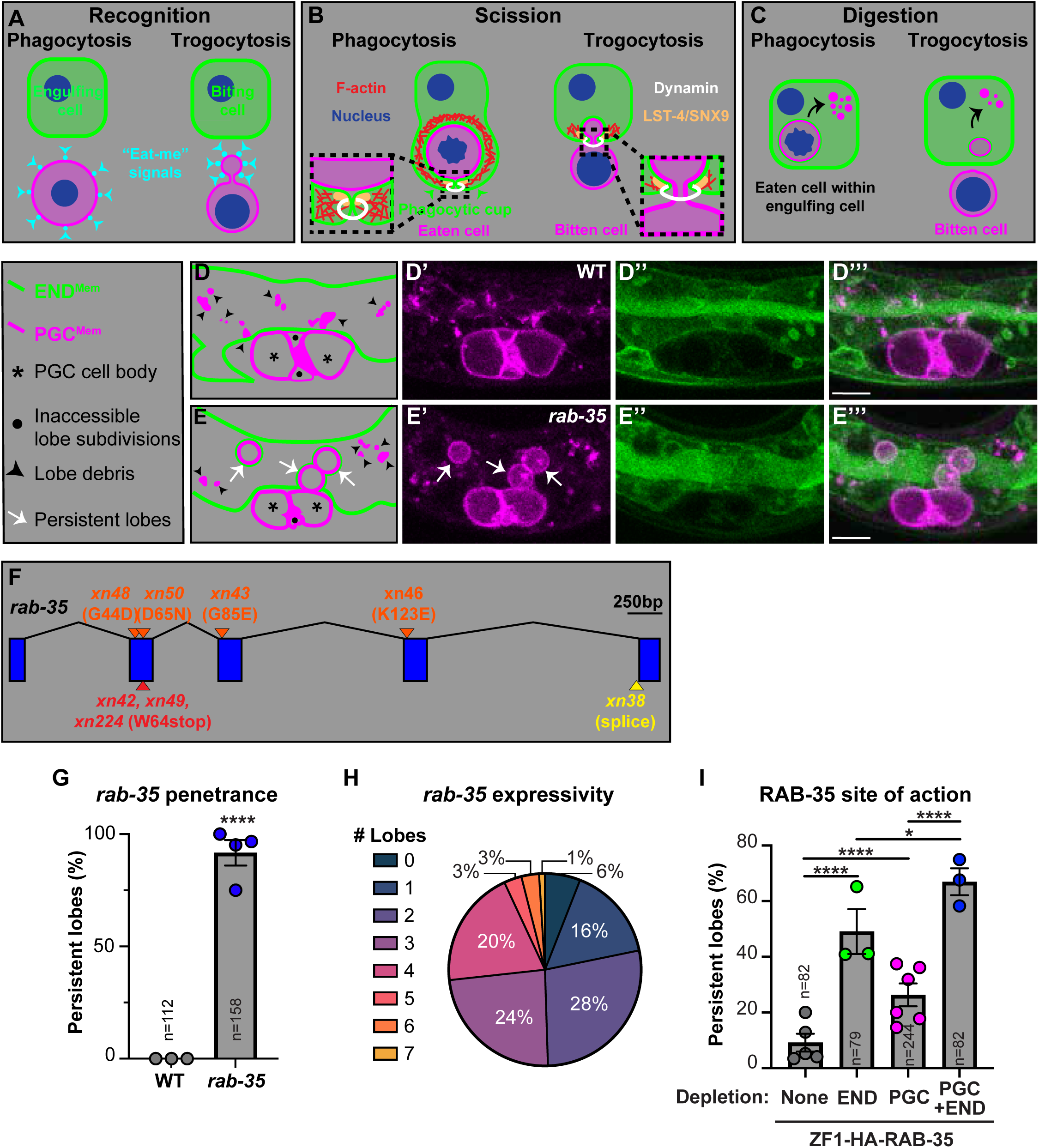
*rab-35* functions in PGCs and endoderm to promote trogocytosis. **(A-C)** Schematic of three steps of trogocytosis (recognition, scission and digestion) and comparisons to phagocytosis. **(D-E’’’)** PGC membranes (PGC^Mem^) and endodermal cell membranes (END^Mem^) in a wild-type L1 larva (D-D’’’) and *rab-35(xn224)* mutant L1 larva (E-E’’’). Schematics of key features are shown in D and E (key on left). Scale bars, 5µm. **(F)** *rab-35* gene structure showing exons (blue boxes), introns (black connecting lines), and positions of mutations (orange, missense; red, nonsense; yellow, splice acceptor). All mutations were isolated in a genetic screen except *xn224*, which is a CRISPR-mediated recreation of the premature stop codon present in *xn42* and *xn49*. **(G)** Penetrance of the persistent lobe phenotype in *rab-35* mutant L1. Circles: percent of L1 with the persistent lobe phenotype in independent replicates (n≥16 animals each replicate); gray box, average of experimental replicates; error bars: SEM; n-values: total number of animals examined across all replicates. ****p-value ≤ 0.0001, Fisher’s exact test. **(H)** Expressivity of the persistent lobe phenotype in *rab-35* mutants, showing the distribution of number of persistent lobes in individual L1 (n=158). **(I)** RAB-35 site of action. The penetrance of the persistent lobe phenotype following depletion of ZF1-HA-RAB-35 in the indicated cells is shown (‘END’, endodermal cells; ‘PGC’, primordial germ cells). See Methods for genotypes used in each depletion. Circles: biological replicates (n≥20 each replicate); error bars: SEM; n-values: total number of animals examined across all replicates. ****p-value ≤ 0.0001, *p-value ≤ 0.05, Fisher’s exact test.

Once internalized, trogocytosed pieces of cells are either recycled or degraded. In the immune system, trogocytosed antigens are commonly recycled to the plasma membrane to elicit an immune response (Baba et al., 2001; Gary et al., 2012; Sabzevari et al., 2001). During development and in specific immune contexts, trogocytosed cargoes are shunted through a degradative pathway within the biting cell that terminates at the lysosomal compartment (Fig. 1C) (Abdu et al., 2016; Gilmartin et al., 2017). Digestion of trogocytosed material has been hypothesized to utilize the same mechanisms that promote digestion of phagocytosed material – a process called phagosome maturation (reviewed in (Pinto and Hengartner, 2012; Wang and Yang, 2016)). Phagosomes mature through sequential interactions with endosomal compartments that are characterized by their association with distinct Rab family small GTPases. These include RAB-35-associated newborn endosomes (Cauvin et al., 2016; Haley and Zhou, 2021), RAB-5-associated early endosomes, and RAB-7-associated late endosomes (reviewed in (Deretic, 2005; Rink et al., 2005)). At each endosomal compartment, phagosomes undergo lipid modifications, which recruit compartment-specific effector proteins that aid in phagosome maturation (Kinchen et al., 2008; Lawe et al., 2000). Finally, late endosomes and lysosomes fuse and cargoes are degraded by hydrolytic enzymes (reviewed in (Kinchen and Ravichandran, 2008)). Trogocytosed material also associates with endosomal and lysosomal compartments during digestion within biting cells (Abdu et al., 2016; Gilmartin et al., 2017), although functional studies demonstrating the importance of phagosome maturation proteins in trogocytosis are lacking.

Here, we examine the mechanisms of a developmentally programmed trogocytic event that occurs when *C. elegans* endodermal cells trogocytose primordial germ cells (PGCs) (Abdu et al., 2016). During mid-embryogenesis, PGCs form an extension, called a lobe, that enriches with organelles before it is bitten off by neighboring endodermal cells (Abdu et al., 2016; Maniscalco et al., 2020; Schwartz et al., 2022). During lobe formation, PGCs polarize such that the nucleus and the lobe are at opposing ends of the cell (Maniscalco et al., 2020). The constriction that produces the lobe and separates it from the cell body forms when a non-mitotic contractile ring partially closes at the lobe neck. Subsequently, lobes bifurcate several times, creating multiple lobe subdivisions. Finally, endodermal cells assemble a scission complex, which includes Rac-induced F-actin, the BAR domain sorting nexin LST-4/SNX9, and DYN-1/Dynamin. The scission complex enriches at the lobe neck within endodermal cells and is required to cut off and internalize the lobe subdivisions (Abdu et al., 2016).

During phagocytosis, dynamin catalyzes membrane fusion by assembling into a helix surrounding two lipid bilayers; helix constriction brings the two membranes close enough for fusion to occur (Fig. 1B) (reviewed in (Antonny et al., 2016; Ramachandran, 2011)). Although dynamin accumulates at the lobe neck within endodermal cells and is essential for lobe scission, structural models suggest that the dynamin helix would be of insufficient diameter to encircle four membrane bilayers (two plasma membrane bilayers from the biting cell and two from the bitten cell) (reviewed in (Antonny et al., 2016; Ramachandran, 2011)). These observations raise the possibility that the bitten cell contributes to trogocytosis by promoting the scission of its own plasma membrane. If so, then dynamin would be required to finish trogocytosis by sealing the endodermal cell membrane around the cut PGC lobe – analogous to its role in the engulfing cell during phagosome sealing.

In this study, we identify *rab-35* as a new trogocytosis regulator with distinct roles in the biting and bitten cell. In biting cells, *rab-35* is required for the digestion of trogocytosed cellular fragments, whereas in the bitten cell, *rab-35* promotes scission. We show that RAB-35 is required for proper phosphatidylinositol 4,5-bisphosphate (PIP_2_) turnover in the biting cell membrane – a process previously shown to be essential for phagosome maturation. Finally, we provide evidence that the ESCRT complex, which can cut membranes from within the cell (reviewed in (Schoneberg et al., 2017)), functions with RAB-35 to regulate trogocytosis. Altogether, our findings reveal *rab-35* as a central regulator of multiple steps in trogocytosis and show that the bitten cell plays an active role in its own trogocytosis.

## RESULTS

### *rab-35* is required for PGC trogocytosis

To better understand how cellular pieces are bitten off and digested, we performed a chemical mutagenesis screen for mutants with defects in PGC lobe trogocytosis. Using a strain co-expressing PGC-specific membrane-targeted mCherry (‘PGC^Mem^’, transgene: *mex-5p::mCherry::PH_PLC∂1_::nos-2 3’ UTR*) and endoderm-specific membrane-targeted GFP (‘END^Mem^’, transgene: *end-1p::GFP::CAAX*), we searched for mutants that retained one or more PGC lobes in L1 larvae – a phenotype we define as ‘persistent lobes.’ In wild type, PGC lobe trogocytosis begins during the middle of embryogenesis (∼2-fold stage) and is completed before the embryo hatches; in newly hatched L1 larvae, only the compact debris of digested PGC lobes within endodermal cells is visible (Fig. 1D-D’’’) (Abdu et al., 2016). Based on the morphometric properties of lobe debris in wild-type L1, we defined persistent lobes in mutant animals as those measuring at least 1.5 µm in diameter, with mCherry enriched on the lobe membrane, and not located at the interface between the two PGCs (where a small number of lobe subdivisions remain in wild type, presumably because they are inaccessible to endodermal cells; Fig. 1D, black dots). Because embryonic PGCs are largely transcriptionally quiescent and must rely on maternally inherited gene products (Schaner et al., 2003), we screened for mutants in the F_3_ generation to leave open the possibility of identifying maternal-effect mutations.

We performed whole genome sequencing to identify the mutations present in nine mutants we isolated that have persistent lobes. Seven mutants contained lesions that alter the coding sequence or splicing of *rab-35*, which is predicted to encode a highly conserved 209 amino acid small GTPase of the Rab family (Fig. 1F, Fig. S1A). Rab GTPases are lipid-modified and associate with membranes (plasma, vesicular, or organellar), where they regulate intracellular trafficking (reviewed in (Homma et al., 2021)). RAB-35 homologues were first identified for their requirement in endocytic recycling, such as the delivery of endosomal compartments to the intracellular bridge in *Drosophila* cells (Kouranti et al., 2006), the transport of T-cell receptors to the immunological synapse (Patino-Lopez et al., 2008), and the recycling of the *C. elegans* yolk receptor back to the oocyte surface (Sato et al., 2008). RAB-35 homologues were subsequently shown to regulate the maturation of newborn endosomes into early endosomes – a function that is important for the digestion of phagocytosed cargoes (Cauvin et al., 2016; Haley et al., 2018).

To confirm that *rab-35* is required for trogocytosis, we engineered a W64stop nonsense mutation, identical to the mutation present in two independently isolated mutants (*xn42, xn49*), into a non-mutagenized background (creating the *rab-35(xn224)* allele) (Fig. 1F). *rab-35(xn224)* is predicted to be a functional null, as it should severely truncate RAB-35 and disrupt its essential GTP-binding domain (Lin et al., 2019) (Fig. S1B). Compared to wild-type L1 larvae, which never displayed persistent lobes (Fig. 1D-D’’’,1G), *rab-35(xn224)* mutant L1 larvae showed a highly penetrant persistent lobe phenotype, with 92% of animals having at least one persistent lobe (Fig. 1E-E’’’,1G). Although we did not test the causality of the other *rab-35* mutant alleles in producing the persistent lobe phenotype, we note that the four missense mutations disrupt highly conserved amino acids (Fig. S1A) and cluster around the GTP binding pocket and effector-binding interface (Fig. S1B) (Lin et al., 2019). Hereafter, we refer to *rab-35(xn224)* animals as *rab-35* mutants.

Each PGC has the potential to produce four or more lobes as a result of lobe bifurcation (Abdu et al., 2016), and *rab-35* mutant L1s contained between 0-7 persistent lobes (2.6 +/- 1.4 S.D.) (Fig. 1E-E’’’,H). Consistent with observations of phagocytosed cell corpses (Haley and Zhou, 2021), we noted a strong temperature dependence of the persistent lobe phenotype of *rab-35* mutants (Fig. S1C), indicating that the process of PGC lobe trogocytosis is sensitive to temperature. Together, these findings reveal that trogocytosis is compromised but not entirely blocked in *rab-35* mutants, suggesting that *rab-35* functions partially redundantly with another pathway to promote PGC lobe trogocytosis.

### RAB-35 functions in both the biting and bitten cell

During phagocytic events, RAB-35 functions within engulfing cells to promote phagosome maturation (Haley et al., 2018; Haley and Zhou, 2021; Kutscher et al., 2018; Wang et al., 2023). To determine whether RAB-35 has an analogous role to digest internalized PGC lobes, we used the ZF1 degron system to deplete RAB-35 protein specifically within endodermal cells. Proteins tagged with the ZF1 (zinc finger 1) degron are degraded rapidly within cells that express the ZIF-1 (zinc finger interacting factor 1) E3 ubiquitin ligase substrate adaptor (DeRenzo et al., 2003; Reese et al., 2000). At the stage of embryogenesis when the PGCs are born, endogenous ZIF-1 activity is restricted to the PGCs (Schwartz et al., 2023), but it can be supplied to specific somatic cells using transgenes (Armenti et al., 2014). We endogenously tagged the N-terminus of *rab-35* with sequences encoding the ZF1 degron and an HA epitope. *rab-35(xn225[zf1::ha::rab-35]); zif-1(-)* mutant animals, which cannot degrade ZF1-HA-RAB-35 because of the *zif-1* mutation, showed only a low penetrance persistent lobe phenotype (Fig. 1I, ‘Depletion: None’). Because *zif-1* mutants with wild-type *rab-35* had no persistent lobe phenotype (0/70 L1 with persistent lobes), we conclude that the low penetrance persistent lobe phenotype of *rab-35(xn225[zf1::ha::rab-35]); zif-1(-)* animals is a result of the ZF1-HA tag, and that ZF1-HA-RAB-35 is largely functional. We were unable to detect ZF1-HA-RAB-35 by immunostaining for the HA epitope.

To test whether RAB-35 functions within the endoderm to promote PGC lobe trogocytosis, we introduced an *elt-2p::zif-1* transgene, which we showed previously promotes degradation of ZF1-tagged proteins specifically in endodermal cells (Armenti et al., 2014), into *rab-35(xn225[zf1::ha::rab-35]); zif-1*(-) mutants. Compared to controls, animals with ZF1-HA-RAB-35 depleted by *elt-2p::zif-1* (hereafter RAB-35^END-^) produced significantly more L1 larvae with persistent lobes (Fig. 1I), indicating that RAB-35 functions at least in part within endodermal cells.

Because RAB-35^END-^ L1 larvae showed a lower penetrance persistent lobe phenotype than *rab-35* mutant L1 larvae (see Fig. 1G), we asked whether RAB-35 is also required in PGCs. For PGC-specific depletion, we examined *rab-35(xn225[zf1::ha::rab-35]); zif-1(+)* animals, taking advantage of endogenous ZIF-1 activity, which we showed previously is specific to PGCs at this stage of embryogenesis (hereafter RAB-35^PGC-^) (Schwartz et al., 2023). Significantly more RAB-35^PGC-^ L1 larvae contained persistent lobes compared to control L1 (Fig. 1I), indicating that RAB-35 also functions within PGCs to regulate trogocytosis. Consistent with a role for RAB-35 in both PGCs and endoderm, the penetrance of the persistent lobe phenotype approached that of *rab-35* mutants when we depleted ZF1-HA-RAB-35 from both cell types simultaneously [*rab-35(xn225[zf1::ha::rab-35]); zif-1(+); elt-2p::zif-1* animals] (Fig. 1I). We conclude that RAB-35 promotes PGC lobe trogocytosis through separable roles in PGCs and endoderm.

### *rab-35* lobes are recognized by endodermal cells but fail during scission and digestion

We next asked which step of trogocytosis *rab-35* promotes – target recognition, scission, or digestion. Target recognition requires that the bitten cell is accessible to, and recognizable by, the biting cell. Although the signals that endodermal cells use to recognize and internalize PGC lobes are not known, endodermal cells appear to be uniquely equipped to internalize PGC lobes, since PGC lobes in endoderm-less mutants persist and remain connected to the PGC cell body (Abdu et al., 2016). To determine if persistent lobes in *rab-35* mutants are properly recognized by endodermal cells, we examined the location of persistent PGC lobes in three-dimensional space relative to the END^Mem^ marker. 99% of *rab-35* persistent lobes were surrounded completely by endodermal cells (e.g. Fig. 1E’’’, n=323 persistent lobes), indicating that nearly all PGC lobes are properly recognized in *rab-35* mutants.

Some persistent lobes in *rab-35* mutants appeared to remain connected to the PGC cell body, whereas others appeared detached. We defined Type I lobes as those located close to the PGC cell body, and when it was possible to discern, attached to the PGC cell body by a thin membrane connection (Fig. 2B,C, orange arrow). These features suggest that Type I lobes have defects in scission and remain connected to the PGC cell body. By contrast, we defined Type II lobes as those present deeper within endodermal cells and lacking a visible membrane connection to the PGC cell body (Fig. 2D,E,F), suggesting that Type II persistent lobes underwent scission but were arrested in digestion. The phenotype of Type II lobes is analogous to that of *rab-35* mutant cell corpses and neuronal exophers, which arrest during the digestion process after they are internalized (Haley et al., 2018; Kutscher et al., 2018; Wang et al., 2023).

**Figure 2.**
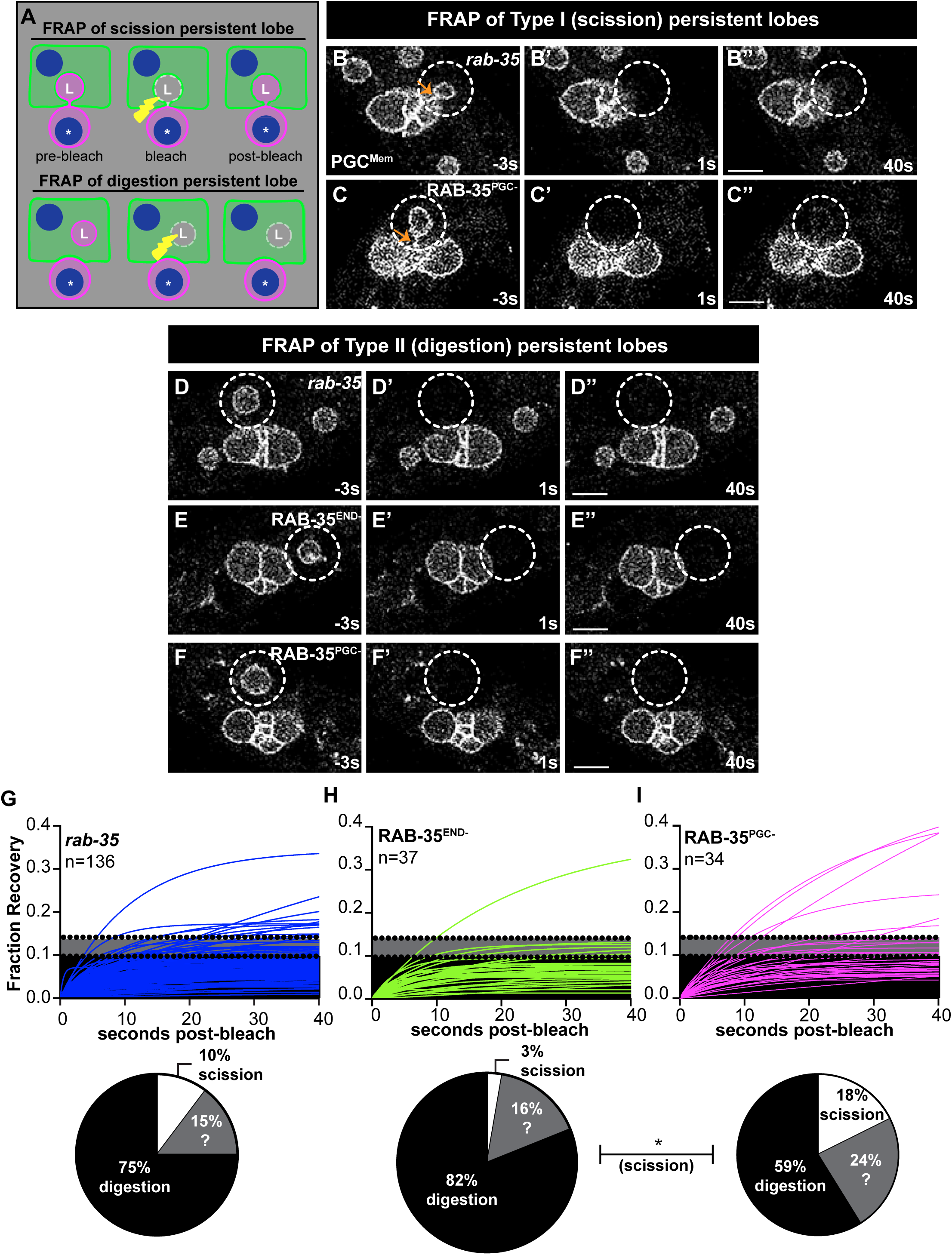
RAB-35 regulates distinct steps of trogocytosis in the biting and target cell. **(A)** Schematic of FRAP assay and expected results for a lobe arrested prior to scission (top row) or digestion (bottom row). PGC^Mem^, magenta; lobe, ‘L’; cell body, asterisk; END^Mem^, green, nucleus, blue; region bleached, yellow lightning bolt and dashed line. **(B-F)** Examples of FRAP experiments showing scission persistent lobes (B-C) or digestion persistent lobes (D-F). Time is relative to photobleaching, photobleached area is surrounded by a dashed circle, genotype indicated. Scale bar, 5µm. **(G-I)** Fluorescence recovery curves, fit to one-phase association models (see Methods) of individual photobleached persistent lobes of indicated genotype. Upper dotted black line: lower threshold for scoring a scission persistent lobe; lower dotted black line: upper threshold for scoring a digestion persistent lobe; area between lines: lobes not classified. Data is summarized in pie charts below. n-values: total number of persistent lobes photobleached across 3 different biological replicates of indicated genotype. *p-value ≤ 0.05, Fisher’s exact test.

To develop quantitative metrics to classify the arrest stage of persistent lobes, we used a fluorescence recovery after photobleaching (FRAP) assay (Abdu et al., 2016). For these experiments, we photobleached the PGC-specific PGC^Mem^ marker on whole individual persistent lobes and plotted fluorescence recovery over time. We expected Type I attached lobes to show continued recovery mediated by diffusion of PGC^Mem^ from the PGC cell body into the lobe (Fig. 2A). By contrast, Type II detached lobes that have completed scission but are arrested in digestion should not recover following photobleaching (Fig. 2A).

We first established classifiers that would distinguish the recovery profiles of attached versus detached persistent lobes. For attached lobes, we plotted the fluorescence recovery curves of photobleached persistent lobes in *lst-4* mutants, which block trogocytosis at the scission step (Abdu et al., 2016). *lst-4* persistent lobes progressively recovered fluorescence over the 40 second imaging period (Fig. S2A), although the extent and rate of recovery varied considerably (which could result from differences in lobe size and width of the thin membrane connection to the cell body). For detached lobes, we examined the recovery curves of a subset of *rab-35* Type II persistent lobes that were located deep within endodermal cells and clearly lacked a membrane connection to the PGC cell body. Type II lobes showed very limited fluorescence recovery, which was restricted to the first few seconds following photobleaching (Fig. S2B); we hypothesize that this limited, rapid recovery results from rapid diffusion of incompletely photobleached mCherry molecules in planes above and below the imaging plane. The fraction recovery of *lst-4* lobes and Type II *rab-35* lobes was significantly different at 40 seconds post-bleach (Fig. S2C). Based on this data set, in all future experiments, we defined attached lobes as those with a fraction recovery greater than any of the detached *rab-35* Type II lobes (> 0.14); detached lobes as those with a lower recovery than any attached *lst-4* lobes (< 0.097), and lobes with a fraction recovery between 0.097 and 0.14 as unclassified.

Applying these classifiers to an unbiased set of photobleached *rab-35* persistent lobes (n = 136), 10% fell into the scission category; 75% into the digestion category, and 15% were unclassified (Fig. 2G). These experiments indicate that *rab-35* is important for both scission and digestion of PGC lobes during trogocytosis.

### RAB-35 regulates distinct steps of trogocytosis within the biting and bitten cell

We next asked whether the percentages of scission versus digestion persistent lobes differ when RAB-35 is depleted in the endoderm versus the PGCs. Only one persistent lobe in RAB-35^END-^ L1 larvae showed the FRAP recovery profile of scission persistent lobes (3%, 1/37 persistent lobes, Fig. 2H), whereas most persistent lobes displayed the recovery profile of digestion persistent lobes (82%, 30/37 persistent lobes, Fig 2H). By contrast, RAB-35^PGC-^ L1 larvae contained a mixture of digestion persistent lobes (59%, 20/34 persistent lobes, Fig. 2I) and scission persistent lobes (18%, 6/34 lobes, Fig. 2I). These data suggest that within endodermal cells, RAB-35 primarily regulates lobe digestion, and within the PGCs, RAB-35 is important for both scission and digestion.

### RAB-35 functions with CNT-1 and ARF-6 to promote trogocytosis

RAB-35 promotes digestion of cell corpses and exophers through the ArfGAP CNT-1/ACAP, which inhibits the small GTPase ARF-6 (Egami et al., 2011; Kutscher et al., 2018; Wang et al., 2023). To test whether CNT-1 and ARF-6 are also important regulators of trogocytosis, we first examined *cnt-1(tm2313)* loss-of-function mutants. *cnt-1* mutant L1 larvae exhibited a strong persistent lobe phenotype (Fig. 3A). Using FRAP to investigate whether *cnt-1* persistent lobes remained connected to PGCs, we observed that 63% remained connected (Fig 3B-B’’, 3D, 19/30 persistent lobes), whereas 26% were cut but not digested (Fig 3C-C’’, 3D 8/30 persistent lobes). Thus, unlike phagocytosis where *cnt-1* only regulates digestion, *cnt-1*, like *rab-35*, is required for both PGC lobe scission and digestion.

**Figure 3.**
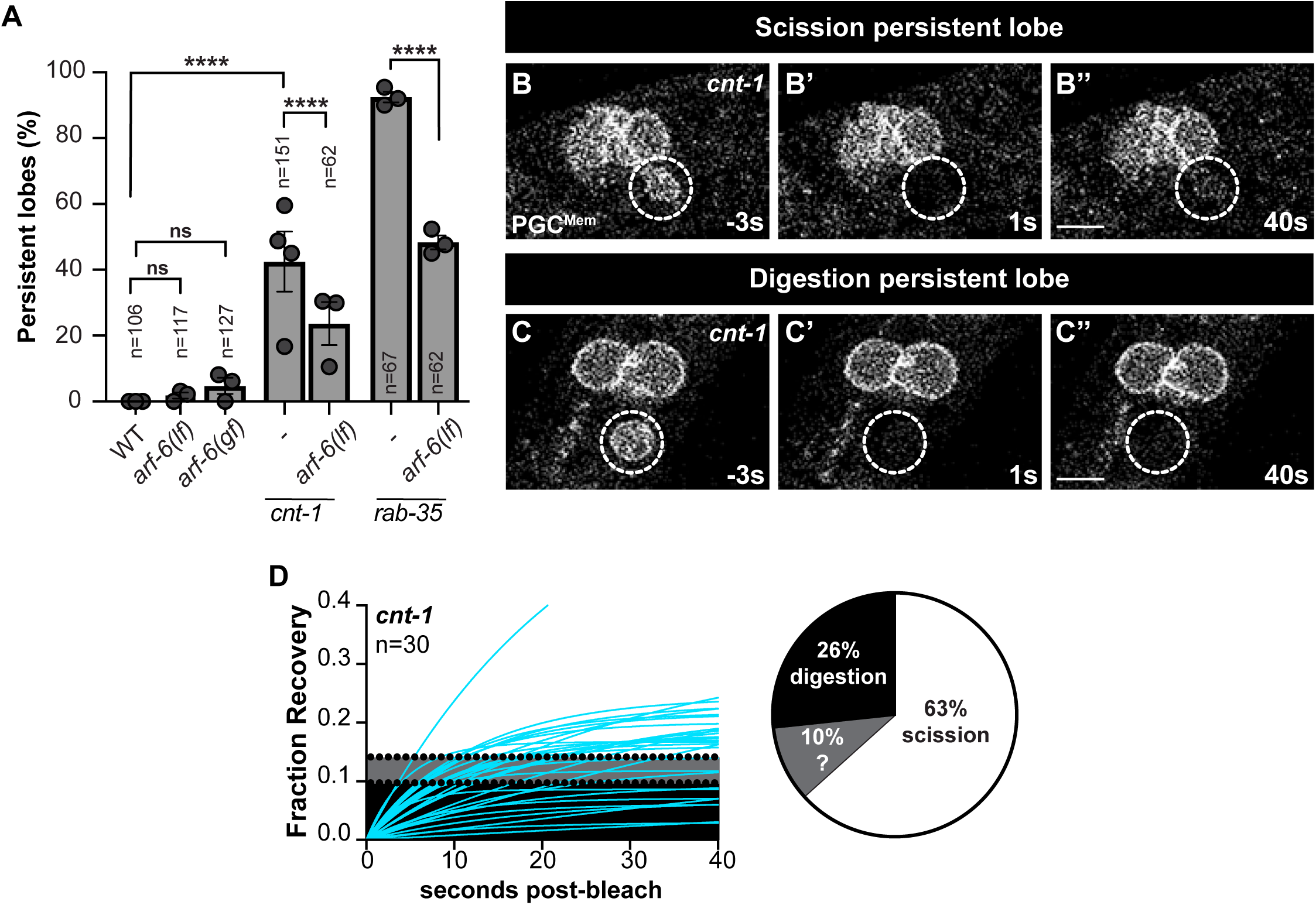
*rab-35* effectors *cnt-1* and *arf-6* regulate PGC lobe trogocytosis. **(A)** Penetrance of persistent lobe phenotype for indicated genotypes. Circles: percent of L1 with the phenotype in independent replicates (n ≥ 19 animals each replicate); gray box, average of experimental replicates; error bars: SEM; n-values: total number of animals examined across all replicates. ****p-value < 0.0001; Fisher’s exact test. **(B-C)** Examples of scission (B) and digestion (C) persistent lobes in *cnt-1* mutant L1. Time is relative to photobleaching; photobleached area is surrounded by a dashed circle, Scale bar, 5µm. **(D)** Fluorescence recovery curves, fit to one-phase association models (see Methods) of individual photobleached persistent lobes in *cnt-1* mutant L1. Upper dotted black line: lower threshold for scoring a scission persistent lobe; lower dotted black line: upper threshold for scoring a digestion persistent lobe; area between lines: lobes not classified. Data is summarized in pie chart. n-values: total number of persistent lobes photobleached across 3 different biological replicates of indicated genotype.

Consistent with observations made during cell corpse and exopher phagocytosis (Kutscher et al., 2018; Wang et al., 2023), the *arf-6(tm1447)* null mutation [‘*arf-6(lf)*’] had no effect on PGC lobe trogocytosis (Fig. 3A). An *arf-6* gain-of-function allele [‘*arf-6(gf)*’] (Kutscher et al., 2018), which was previously shown to mimic the *cnt-1* and *rab-35* neuronal exopher and cell corpse digestion defects, caused only a weakly penetrant persistent lobe phenotype that was not significantly different from wild type (Fig. 3A). However, consistent with RAB-35 and CNT-1 inhibiting ARF-6 to regulate trogocytosis, *arf-6(lf)* partially suppressed the persistent lobe phenotype of both *cnt-1* and *rab-35* mutants (Fig. 3A). All together, these data suggest that RAB-35 and CNT-1 regulate trogocytosis in part by inhibiting ARF-6, and in part through a parallel, ARF-6-independent pathway.

### RAB-35 localization in endodermal cells and PGCs

Because RAB-35 enriches around membranes prior to the appearance of early endosomal marker RAB-5 during the phagocytosis of cell corpses (Haley and Zhou, 2021), we predicted that RAB-35 would enrich around freshly cut PGC lobes within endodermal cells during trogocytosis. To establish where endogenous RAB-35 localizes in biting and bitten cells, we tagged the *rab-35* gene with two sequences encoding the 11^th^ ß-barrel of GFP and used transgenes to express the complementing GFP_1-10_ fragment in either somatic cells or PGCs. Because only the reconstituted GFP_1_-_10_ + GFP_11_ protein fluoresces, this approach allowed us to distinguish between RAB-35 localization patterns in the two cell types (Cabantous et al., 2005; Kamiyama et al., 2016).

Reconstituted somatic GFP_11_-RAB-35 (hereafter GFP_11_-RAB-35^SOMA^) was broadly expressed, and its punctate subcellular localization was consistent with endomembrane localization, as previously reported for transgenic RAB-35 (Sato et al., 2008). GFP_11_-RAB-35^SOMA^ loses some functionality within this genetic background, as animals displayed a significant persistent lobe phenotype when somatic GFP_1_-_10_ was present (Fig. S3A), although the phenotype was much less severe than that of *rab-35* null mutants.

To determine if, and when, GFP_11_-RAB-35^SOMA^ enriches around PGC lobes, we examined lobes in wild-type animals during the embryonic stage when they are bitten from PGCs. Because the relative timing of trogocytosis steps is variable (Abdu et al., 2016), we compared GFP_11_-RAB-35^SOMA^ localization around lobes that remained in contact with the PGC cell body, which based on our FRAP experiments we inferred were predominantly still connected and had not yet undergone scission, to non-contacting lobes, which we presumed were recently bitten off and in the process of digestion (Fig. 4A). Although contacted lobes displayed some enrichment of GFP_11_-RAB-35^SOMA^ around their surfaces, we noted a significantly stronger enrichment around non-contacted lobes (Fig. 4B-D). Since GFP_11_-RAB-35^SOMA^ is not visible within the PGCs in this background (Fig 4B’-B’’, 4C’-C’’), and because lobes are embedded within endodermal cells, this somatic pool of GFP_11_-RAB-35^SOMA^ is within endodermal cells. These data suggest that RAB-35 becomes more enriched on PGC lobes after scission has occurred. This localization pattern is analogous to that described for RAB-35 in phagocytic events and is consistent with our finding that RAB-35^END^ promotes PGC lobe digestion.

**Figure 4.**
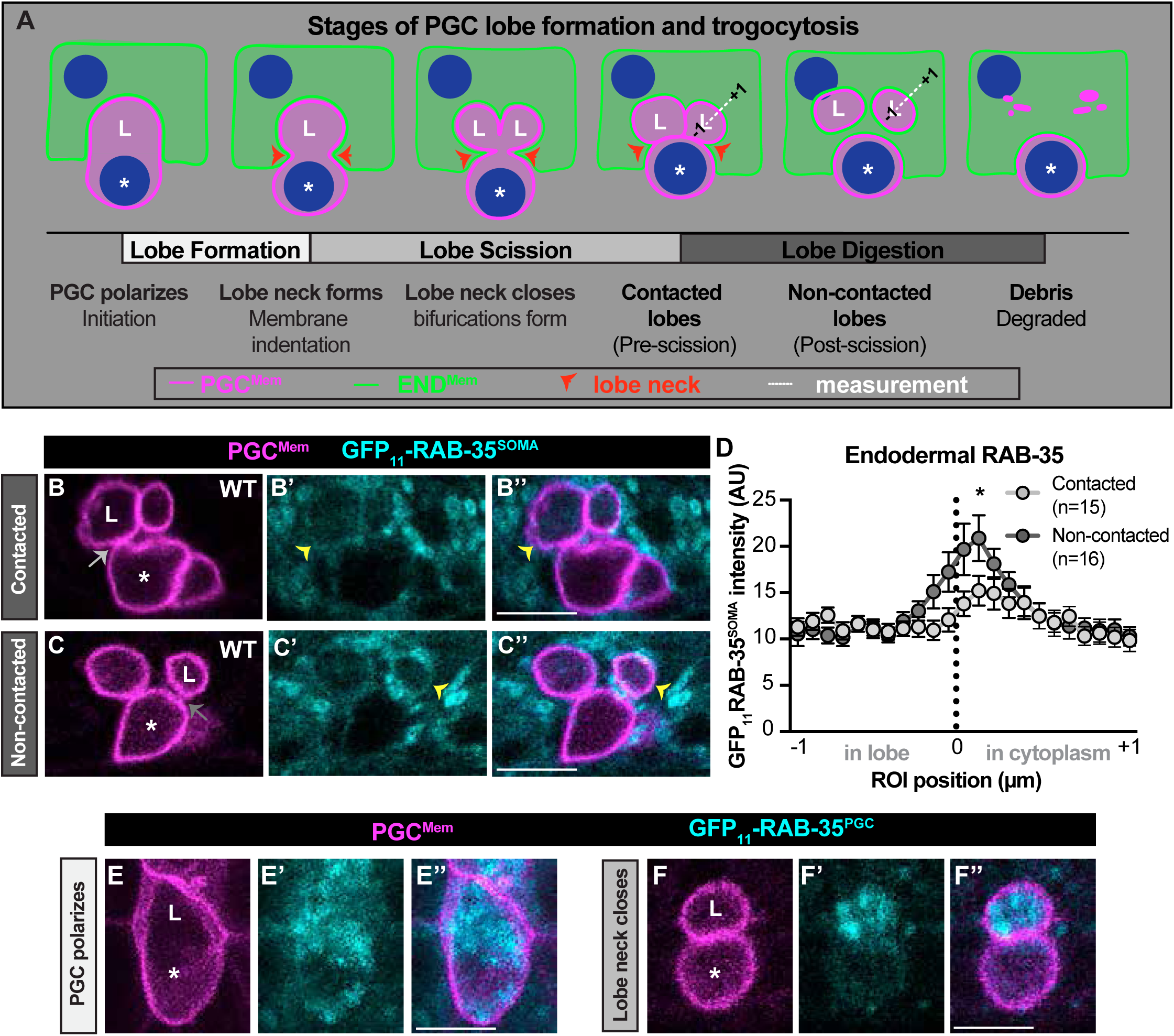
RAB-35 localization pattern in PGCs and endoderm. **(A)** Schematic of the stages of PGC lobe formation and trogocytosis. **(B-C)** Micrographs (single plane) of contacted and non-contacted lobes. Lobe, ‘L’; cell body, asterisk; arrow, contact (lighter gray) or space between the lobe and the PGC cell body (darker gray); GFP_11_-RAB-35^SOMA^ surrounding the lobe, yellow arrowheads. Scale bars, 5µm. **(D)** Quantification of fluorescence intensity of GFP_11_-RAB-35^SOMA^ at non-contacted lobes (light gray circles) and contacted lobes (dark gray circles). 1 µm ROI crossing lobe boundary schematized (A), with 0 position at lobe boundary, negative positions in lobe, and positive positions in endodermal cell cytoplasm. Circles: average fluorescence intensity; error: SEM; n-values: total number of lobes within each category analyzed across 4 biological replicates. *p-value ≤ 0.05, multiple unpaired t-tests. **(E-F)** Micrographs (E) z-stack (F) single plane of GFP_11_-RAB-35^PGC^ (cyan) when PGC is polarizing to form a lobe (E) and after closure of the lobe neck (F). Lobe, ‘L’; cell body, asterisk. Scale bars, 5µm.

We next examined GFP_11_-RAB-35 localization using a transgene that expresses GFP_1-10_ specifically in PGCs (*mex-5p*:*gfp1-10*::*nos-2* 3’UTR) (Schwartz et al., 2023). In contrast to GFP_11_-RAB-35^SOMA^, GFP_11_-RAB-35 co-expressed with PGC-specific GFP_1-10_ (hereafter GFP_11_-RAB-35^PGC^) did not cause a persistent lobe phenotype (Fig. S3A). Consistent with our observations in endodermal cells, and with previous observations in other cell types (Patino-Lopez et al., 2008; Sato et al., 2008), GFP_11_-RAB-35^PGC^ was found throughout the PGCs and localized in a pattern suggesting endomembrane association (Fig. 4E-4F). We did not note a strong enrichment of GFP_11_-RAB-35^PGC^ at the lobe neck, either during lobe initiation (Fig. 4E) or as the lobe neck closes (Fig. 4F), suggesting either that RAB-35 regulates scission from sites distant to the lobe neck, or that a subpopulation of RAB-35 at the lobe neck is important for scission.

### RAB-35 regulates PIP_2_ loss on lobe membranes within endodermal cells

During phagocytosis, RAB-35 inhibits ARF-6 to promote the loss of PIP_2_ on the engulfing cell membrane surrounding phagocytosed cargo (Kutscher et al., 2018; Wang et al., 2023). PIP_2_ conversion is thought to be a key step in phagosome maturation and has been proposed as the reason engulfed cell corpses persist in *rab-35* mutants (Haley and Zhou, 2021; Kutscher et al., 2018). The enrichment of GFP_11_-RAB-35 surrounding recently trogocytosed PGC lobes suggests that RAB-35-mediated PIP_2_ conversion might also be necessary to degrade trogocytosed cellular pieces. We first examined the localization of PIP_2_ in endodermal cells by expressing mCherry fused to the PIP_2_-binding PH domain of rat PLC∂1 (hereafter PIP_2_^END^). Because PGC lobe trogocytosis occurs when the embryo is moving rapidly within the eggshell (complicating live imaging), we examined PIP_2_^END^ at single time points, comparing its enrichment surrounding lobes that contacted the PGC cell body (inferred to be largely pre-scission) to non-contacted lobes that had not yet condensed into debris (inferred to be post-scission). For quantification, we measured absolute levels of the PIP_2_^END^ reporter in a 2 µm ROI line spanning the lobe membrane, as schematized in Fig. 4A. On contacted lobes, PIP_2_^END^ signal peaked adjacent to the PGC membrane in both wild-type and *rab-35* mutants (Fig. 5A-B’’’), although the amount of PIP_2_^END^ was significantly higher overall in *rab-35* mutants compared to wild type (Fig. 5E). In presumed cut lobes (non-contacted), PIP_2_^END^ surrounding PGC lobe membranes largely disappeared in wild type (Fig. 5C-C’’’,F), similar to observations made during phagocytosis. By contrast, in *rab-35* mutants, PIP_2_^END^ remained at high levels surrounding lobes that appeared to have completed scission (Fig. 5D-D’’’, 5F). Together with our observation that GFP_11_-RAB-35^SOMA^ enriches on lobes after scission has occurred, these findings suggest that RAB-35-mediated PIP_2_ loss on the endodermal membrane surrounding cut PGC lobes is a key step in their degradation.

**Figure 5.**
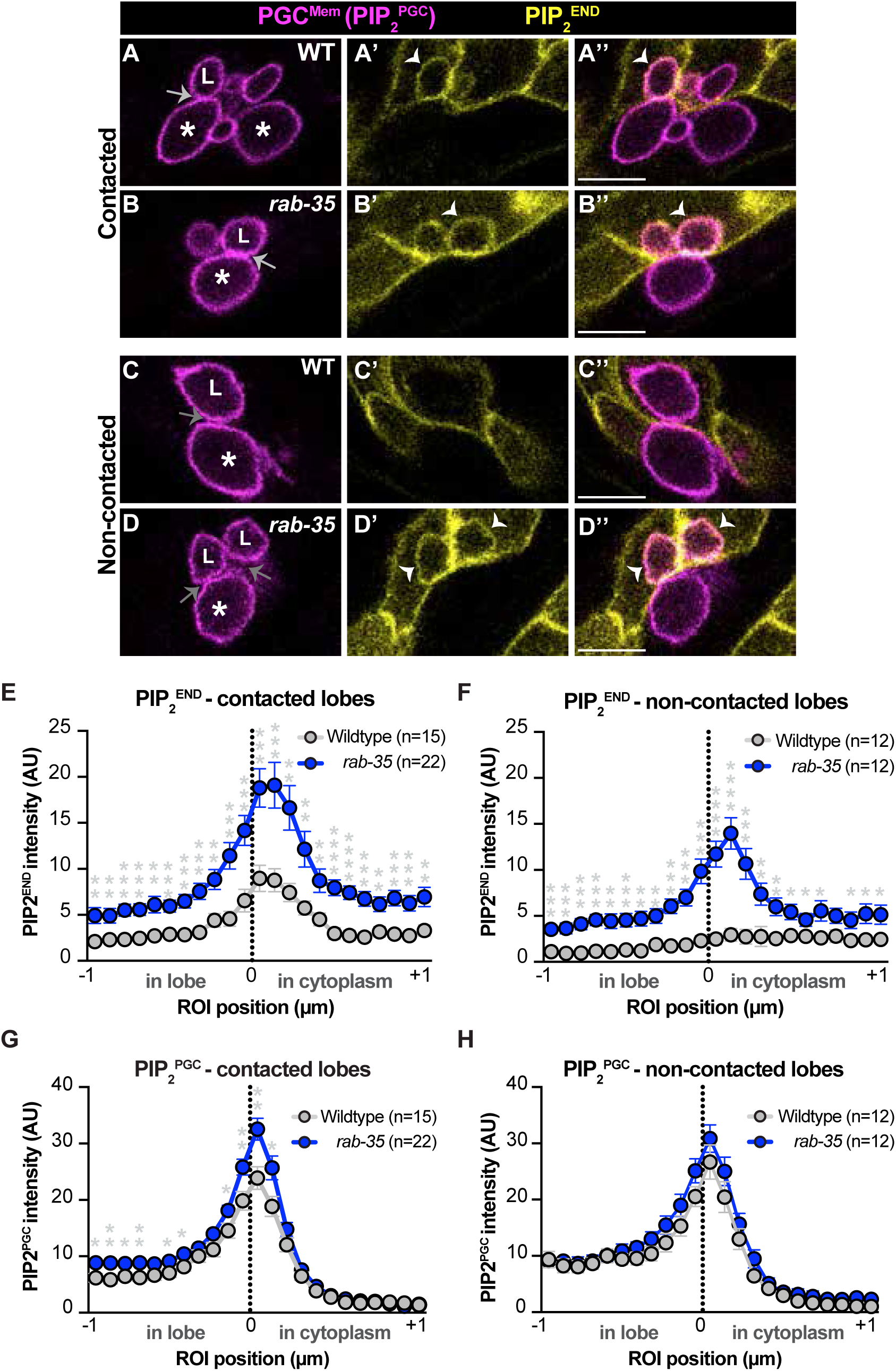
RAB-35 regulates PIP_2_ loss on lobe membranes within endodermal cells. **(A-D)** Micrographs of reporters for PGC and endodermal PIP_2_ (recognized by the PH domain of rat PLC∂1). Arrow, contact (lighter gray) or space between the lobe and the PGC cell body (darker gray); white arrowhead, PIP_2_^END^ surrounding PGC lobe membrane. Scale bars, 5µm. **(E-H)** Quantification of fluorescence intensity of PIP_2_ reporters in contacted and non-contacted lobes, as indicated. ROIs were drawn as depicted in 4A, with positions -1 µm – 0 µm: within the lobe, position 0µm at the PGC lobe membrane (black, vertical dotted line), and positions 0 µm – +1 µm within the endodermal cytoplasm. Circles, average intensity across three biological replicates; error, SEM; n-values, total number of lobes analyzed. Statistics: multiple, unpaired t-tests, *p-value ≤ 0.05, **p-value ≤ 0.01, ***p-value ≤ 0.001, ****p-value ≤ 0.0001

Because we noted overall higher PIP_2_^END^ levels in *rab-35* mutant endodermal cells, we also examined PIP_2_ levels in PGCs using the PGC^Mem^ marker, which contains the PIP_2_-binding PLC∂1 PH domain (hereafter PGC^PIP2^). Contacted (presumed attached) lobes had significantly higher PIP_2_^PGC^ signal in r*ab-35* mutants compared to wild type (Fig. 5G), although this difference was lost in non-contacted (presumed cut) lobes that appeared to have completed scission (Fig. 5H). These data raise the possibility that aberrant PIP_2_ regulation in *rab-35* mutant PGCs interferes with key steps of trogocytosis, including lobe formation or lobe scission.

### The ESCRT complex regulates lobe scission

How does RAB-35 promote the scission step of trogocytosis within the bitten cell? In mammalian cells, RAB35 regulates cytokinesis (Kouranti et al., 2006), raising the possibility that PGC lobe trogocytosis defects are a secondary consequence of slowed or delayed non-mitotic contractile ring closure during PGC lobe formation. To address this possibility, we first asked whether the events preceding scission, including PGC polarization and contractile ring formation (Fig. 4A), are perturbed in RAB-35^PGC-^mutants. Using the migration of the nucleus as an indicator of PGC polarization, and PGC plasma membrane indentation as an indicator of contractile ring function (see Fig. 4A), we observed that PGCs polarized and began to constrict their contractile ring at similar developmental stages in RAB-35^PGC-^ animals as in wild type (Fig. S4A-B). These findings strongly suggest that scission defects in RAB-35^PGC-^ mutants are unlikely to result indirectly from improper lobe formation.

During late cytokinesis, mammalian RAB35 promotes the removal of PIP_2_ at the contractile ring (Dambournet et al., 2011) to allow abscission (Kumar et al., 2023). The <u>E</u>ndosomal <u>S</u>orting <u>C</u>omplexes <u>R</u>equired for <u>T</u>ransport (ESCRT) complex, which assembles into rings at the site of future membrane scission, mediates abscission and other membrane fission events defined as having reverse topology (cut by pulling membranes together from the inside, as opposed to squeezing together from the outside) (reviewed in (Schoneberg et al., 2017)). To determine if RAB-35 promotes the cutting of PGC lobes from within the cell, we performed experiments to determine if the ESCRT complex contributes to PGC lobe scission. *C. elegans* has a highly conserved suite of ESCRT proteins composed of five classes (ESCRTs -0, -I, -II, -III, and VPS4/VPS-4) (reviewed in (Henne et al., 2011; Hurley, 2015)). We first examined whether ESCRT was expressed in PGCs. Using a split GFP approach, we endogenously tagged *tsg-101* (ESCRT-I) with sequences encoding GFP_11_ (*tsg-101*(*xn237*[*tsg-101::2xgfp_11_*]) and introduced PGC-specific GFP_1-10_, as described above (hereafter TSG-101^PGC^). We observed no persistent lobe phenotype within this genetic background, indicating that TSG-101-GFP_11_ is functional (Fig. S3B). In wild-type PGCs, TSG-101^PGC^ was present diffusely throughout the cell, and concentrated in brighter puncta within lobes, including just prior to lobe scission (Fig. 6A). TSG-101^PGC^ showed a similar localization in *rab-35* mutant PGCs (Fig. 6B, S5A-C).

**Figure 6.**
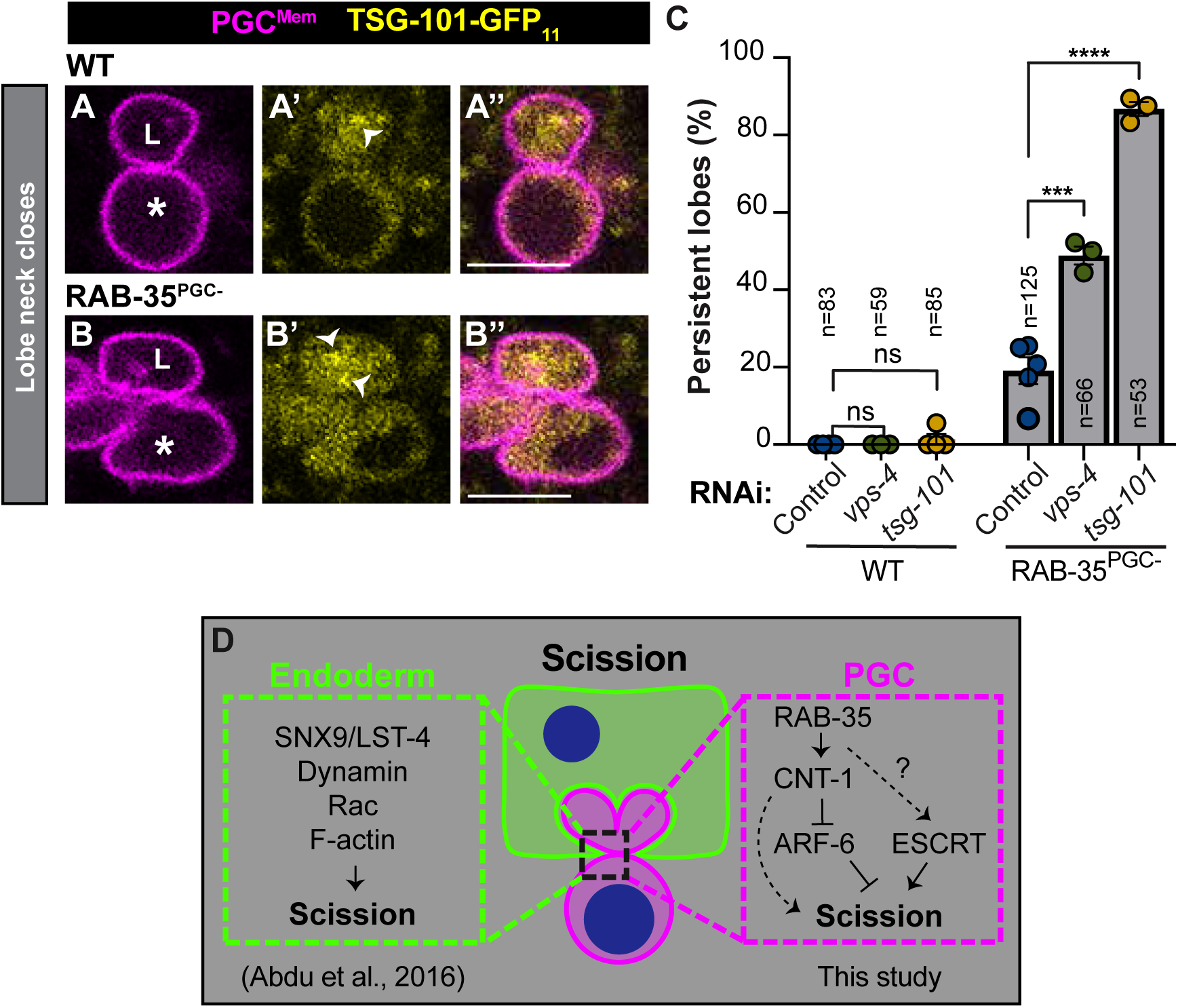
The ESCRT complex and RAB-35^PGC^ regulate trogocytosis through parallel pathways. **(A-B)** Micrographs of TSG-101^PGC^ within wild-type (A-A’’) and RAB-35^PGC-^ (B-B’’) PGCs. Arrowheads, TSG-101 puncta identified by Fiji 3D object counter (see Methods). **(C)** Penetrance of persistent lobe phenotype after RNAi-mediated knockdown in the indicated genetic background (WT or RAB-35^PGC-^). Circles, percent persistent lobes within a single biological replicate (n ≥ 15 animals); bars, average penetrance of all replicates; n-values: number of lobes analyzed across ≥ 3 biological replicates. Error bars: SEM, statistics: Fisher’s exact test, ns p-value > 0.05, ***p-value ≤0.001, ****p-value ≤ 0.0001. **(D)** Model of PGC scission. Endoderm scission complex, identified in (Abdu et al., 2016), summarized in green dotted box. PGC scission complex, described here, summarized in pink dotted box. Unknown regulators, question marks; positive genetic interactions, arrows; lines with bars, inhibitory genetic interactions; dashed lines, uncertain relationships.

Next, we examined ESCRT function in trogocytosis. ESCRT function is essential for embryonic development (Roudier et al., 2005), precluding the use of mutants to examine PGC lobe trogocytosis. Therefore, we used RNAi to partially knock down levels of ESCRT proteins, asking if partial depletion enhanced the trogocytosis defects of RAB-35^PGC-^ mutants. We examined two ESCRT genes: (1) the ESCRT-I component *tsg-101*, (Carlton and Martin-Serrano, 2007; Morita et al., 2007); and (2) the AAA-ATPase *vps-4*, which functions downstream of TSG-101 and is thought to have a direct role in membrane scission (Elia et al., 2011; Schoneberg et al., 2018). Within a wild-type background, partially knocking down *tsg-101* produced only a very low penetrance persistent lobe phenotype, whereas *vps-4* partial knockdown had no effect on PGC lobe trogocytosis (Fig. 6C). However, *vps-4* or *tsg-101* partial knockdown significantly enhanced the persistent lobe phenotype of RAB-35^PGC-^ L1 larvae (Fig. 6C). Thus ESCRT, like RAB-35, promotes trogocytosis of PGC lobes.

## DISCUSSION

Here, we characterize a new regulator of trogocytosis – RAB-35 – which regulates distinct steps of trogocytosis in the biting and bitten cells. Our analysis of RAB-35 in biting (endodermal) cells shows that it is important for digesting trogocytosed PGC lobes. In the bitten cell (PGCs), RAB-35 promotes scission in conjunction with the ESCRT membrane scission complex. We found that RAB-35 works together with downstream effectors CNT-1 and ARF-6, which are known to control levels of the phospholipid PIP_2_, and that *rab-35* mutants have increased PIP_2_ levels, suggesting that improper PIP_2_ regulation could be responsible for trogocytosis defects in *rab-35* mutants. Our findings provide evidence that membrane scission during trogocytosis is not fueled solely by the biting cell - the bitten cell also promotes scission of its own plasma membrane.

### A common pathway for digesting phagocytosed or trogocytosed cargo

We showed previously that late endosomal marker RAB-7 and lysosomal protein LMP-1 associate with trogocytosed PGC lobes as they are being digested (Abdu et al., 2016). RAB-7 and LMP-1 are known to promote phagosome maturation, suggesting that cells or cell fragments taken up by phagocytosis or trogocytosis might be digested through a common pathway. The requirement of RAB-35 in endodermal cells to promote digestion of PGC lobes provides functional evidence for this hypothesis, as *rab-35* was shown previously to promote the digestion of phagocytosed cell corpses and neuronal exophers (Haley et al., 2018; Kutscher et al., 2018; Wang et al., 2023).

During phagocytosis, RAB-35 localizes around internalized cell corpses and acts through CNT-1 and ARF-6 to promote turnover of PIP_2_, which is a prerequisite to phosphatidylinositol 3-phosphate (PI(3)P) accumulation on the engulfing cell membrane. In turn, PI(3)P recruits downstream effectors important for phagosome maturation and digestion, including early endosomal antigen 1 (EEA-1) (Lawe et al., 2000), and Rabenosyn-5 (RABS-5) (Nielsen et al., 2000), which localize through PI(3)P-recognition domains. We showed that RAB-35 enriches around trogocytosed PGC lobes, functions with CNT-1 and ARF-6, and is required for the turnover of PIP_2_ on PGC lobe trogosomes. These commonalities suggest that the molecular function of RAB-35 in promoting digestion of phagocytosed cells and trogocytosed cellular pieces is conserved.

### RAB-35 functions in the bitten cell to promote scission during trogocytosis

Because previous studies implicated *rab-35* function in promoting the digestion of phagosomes, we were surprised to discover that *rab-35* functions not only in the digestion of trogocytosed PGC lobes but also in their scission. Moreover, we showed that removing RAB-35 specifically from PGCs results in lobe scission defects, indicating that RAB-35 contributes to distinct steps of trogocytosis in the biting and bitten cell. An important implication of these findings is that scission during trogocytosis is not accomplished solely through actions of the biting cell. Rather, the bitten cell actively participates in its own scission.

Because RAB-35 has no known or predicted membrane bending or severing activity, it very likely functions to promote scission indirectly. Our genetic enhancement experiments show that the ESCRT complex, which is known to sever membranes from within the cell by pulling them together, also contributes to scission together with *rab-35*. Although the relationship between RAB-35 and ESCRT remains unclear, we noted no difference in the amount of ESCRT complex protein TSG-101 in the RAB-35^PGC-^background. It is possible that RAB-35 is needed for ESCRT activity, or alternatively, that ESCRT and RAB-35 function in parallel to promote scission. To our knowledge, there is no known molecular connection between homologues of RAB-35 and the ESCRT complex in other systems. However, during mammalian cytokinesis, Rab35 and ESCRT both promote abscission to separate the two daughter cells (Carlton and Martin-Serrano, 2007; Kouranti et al., 2006; Morita et al., 2007), and loss of Rab35 results in abnormally high levels of PIP_2_ at the cleavage furrow (Dambournet et al., 2011). It is possible that the increased PIP_2_ levels we observed in PGCs of *rab-35* mutants similarly interferes with ESCRT function during the scission step of trogocytosis.

We found that removing RAB-35 from PGCs also results in some persistent lobes that complete scission but arrest in digestion. One possible explanation of this phenotype is that some *rab-35* mutant lobes undergo delayed scission and are internalized just prior to our FRAP experiments. An argument against this model is that PGC lobes in wild-type embryos are rapidly digested into compact debris after scission is complete (Abdu et al., 2016), although it remains possible that digestion occurs less rapidly at stages after trogocytosis is normally complete. Another possibility is that RAB-35 regulates proteins or phospholipids in the PGC lobe membrane that promote digestion indirectly, for example by signaling to surrounding endodermal cell membranes.

### Coordination of scission between the biting and bitten cells

Previously, we showed that scission of PGC lobes is mediated by a scission complex, which includes LST-4/SNX9, the membrane-severing protein dynamin, and F-actin, that accumulates on the endodermal face of the PGC lobe neck. In *lst-4* mutants, PGC lobes cannot undergo scission, and restoring *lst-4* function specifically within endodermal cells rescues this phenotype (Abdu et al., 2016). However, here we have shown that PGCs use RAB-35 to promote the scission of their own membrane during trogocytosis. Together, these findings indicate that scission during trogocytosis requires activities in both the biting and bitten cell, implying that these events must be coordinated for scission to occur. Communication between endodermal cells and PGCs could involve active signaling from proteins present on either side of the lobe neck. Alternatively, constriction at the lobe neck within one cell type might be required for the localization or activity of scission proteins within the other cell type. Collectively, our findings suggest that therapeutic interventions to modify the efficiency of trogocytosis could be designed to target distinct pathways in either the biting or bitten cell.

## ACKNOWLEDGEMENTS

We thank Shai Shaham for providing worm strains; Michael Cammer for assistance with imaging and image analysis; Exon-Intron Graphic Maker (Nikhil Bhatla, wormweb.org/exonintron) for assistance in creating gene structures; Ann Wehman, Holger Knaut, and Barth Grant for helpful discussion, and Nance lab members for comments on the manuscript. Some strains were obtained from the *Caenorhabditis* Genetics Center, which is funded by NIH Office of Research Infrastructure Programs (P40 OD010440). Some experiments were performed in conjunction with the NYU Langone Microscopy Laboratory (RRID: SCR_017934) and the Genome Technology Center (RRID: SCR_017929). Funding for this study was provided by the National Institutes of Health through research grant R35GM118081 (J.N.) and training grant T32HD007520-24 (J.M.), and by a Howard Hughes Medical Institute International Student Research fellowship (Y.A.).

## METHODS

### Worm culture and strains

Worms were maintained as previously described (Brenner, 1974). Worms were grown at ∼23°C and shifted to 25°C ∼72 hours prior to experiments, unless noted otherwise. Table S1 includes a list of strains used in this study.

### Genetic screen and mutant identification

*rab-35* mutants were isolated in a previously described non-clonal, maternal-effect mutagenesis screen for L1 larvae with persistent PGC lobes (Abdu et al., 2016). To identify causal mutations, some mutant strains (FT1406, FT1463) were first backcrossed to the starting non-mutagenized strain (FT2523) and F2 progeny were singled and allowed to self-fertilize, establishing recombinant lines (n ≥ 9 recombinant lines). Individual F2 recombinants were classified as mutant or non-mutant by analyzing the phenotype of their progeny. For these mutant strains, mutant and non-mutant recombinant lines were pooled separately; for the remaining mutant strains (FT1464, FT1469, FT1504, FT1542, FT1543), lines were propagated without backcross. Genomic DNA (gDNA) was isolated as described (Abdu et al., 2016) and submitted to the NYU Genome Technology Center for library preparation and whole genome sequencing (Meyer et al., 2026). High and moderate impact variants were identified through the sibling subtraction method (Joseph et al., 2018). Seven of nine sequenced mutant strains contained mutations predicted to alter the protein coding sequence (missense, nonsense, or splice site mutations) of *rab-35*. To confirm that *rab-35* disruption causes a persistent lobe phenotype, the premature stop codon present within *rab-35(xn42)* and *rab-35(xn49)* was introduced into a non-mutagenized background using CRISPR genome editing. The resulting mutant, *rab-35(xn224[rab-35W64stop])*, showed a persistent lobe phenotype (see Fig. 1G).

### Mutant phenotype quantification

Individual L1 were screened for a persistent lobe phenotype on a Zeiss AxioImager with 63X 1.4NA oil objective or a Leica laser scanning confocal (SP8) with 63X 1.4NA oil objective. L1 larvae were scored as mutant if they contained at least one persistent lobe, which we defined as >1.5 µm in diameter with PGC^Mem^ enriched at the surface. Mutant penetrance was defined as the percent of mutant L1 larvae that contained one or more persistent lobes. Mutant expressivity was defined as the number of persistent lobes within a mutant L1.

### CRISPR gene editing

Endogenously tagged alleles engineered in this study were created using CRISPR/Cas9 homology-directed repair, as described (Paix et al., 2015). tracrRNA, custom crRNAs, and repair template oligonucleotides were synthesized by Integrated DNA Technologies (IDT), and purified Cas9 protein was obtained from the UC-Berkeley QB3 MacroLab. Repair oligos included ∼35bp homology arms. When necessary, silent mutations were introduced into the PAM site or crRNA hybridization region to prevent re-cutting. Candidate edits were confirmed by PCR and sequencing. crRNA and repair template sequences are listed in Table S2.

### Tissue-specific RAB-35 degradation

RAB-35 was conditionally degraded from PGCs or endoderm using the ZF1/ZIF-1 degron strategy (Armenti et al., 2014). Using CRISPR/Cas9, sequences encoding the ZF1 zinc finger degron and an HA epitope tag were inserted immediately downstream of the ATG at the 5’ end of the *rab-35* gene, creating *rab-35(xn225[zf1::ha::rab-35])*. ZF1-HA-RAB-35 was degraded specifically from PGCs using endogenous ZIF-1 activity, which during mid-embryogenesis is restricted to PGCs (Schwartz et al., 2023) [genotype: *rab-35(xn225[zf1::ha::rab-35]); zif-1(+)*]. ZF1-HA-RAB-35 was degraded specifically from endodermal cells by eliminating endogenous ZIF-1 activity using the *zif-1(gk117)* mutation and supplying ZIF-1 activity specifically to endodermal cells using an *elt-2p*:*zif-1* transgene, which we described previously (Armenti et al., 2014) (genotype: *rab-35(xn225[zf1::ha::rab-35]); zif-1(gk117); xnEx561 [elt-2p::zif-1 + unc-122p::gfp]*). ZF1-HA-RAB-35 was degraded from both PGCs and endoderm using genetic background *rab-35(xn225[zf1::ha::rab-35]); zif-1(+); xnEx561 [elt-2p::zif-1 + unc-122p::gfp]*.

### FRAP experiments

Gravid mothers were dissected in M9 buffer to release embryos, which were allowed to hatch overnight at 25°C. L1 larvae were mounted on 10% agarose pads and paralyzed in ∼5mM levamisole. FRAP was performed on a laser scanning confocal microscope (Leica SP8), using a 63X 1.4NA oil or water objective with an Argon tunable laser, 488/561nm lasers and a PMT detector. Regions of interest (ROIs) were drawn around persistent lobes (avoiding PGC cell body signal). Three pre-bleach time points, one time point just after bleach, and 40 post-bleach time points were collected at 1 second time intervals.

To generate recovery curves, regions of interests (ROIs) were drawn around persistent lobes (avoiding PGC cell body signal) using Fiji (ImageJ), and mean intensities (F_rawData_) were calculated across time points. FRAP data normalization was modified from suggested methods (Kang et al., 2015). To control for background signal, we calculated the mean intensity value of the background using a ROI of the same size, generating F_background_. F_background_ was subtracted from both F_rawData_ and the mean intensity value of the whole image, F_rawWhole_, generating F_data_ and F_whole_. To control for photobleaching, we divided F_whole_ from F_data_ generating F_timepoint_ for each time point. The three F_timepoint_ pre-bleach time points were averaged and served as F_pre-bleach_. F_bleach_ and F_post-bleach_ were divided by F_pre-bleach_ to determine fraction recovery and plotted over time. F_post-bleach_ was then scaled from 0-1 by subtracting the F_bleach_ value. Fraction recovery curves were then plotted using XY, one phase association, exponential models on GraphPad/Prism – bleach point fixed at y=0. Values at 40-seconds post-bleach were calculated using the equation of computed exponential curves. To establish criteria for classifying scission versus digestion persistent lobe recovery thresholds, the ranges of all *lst-4* persistent lobes (n=38), and a randomly generated subset of *rab-35* Type II persistent lobes (n=38) were assessed.

### RNAi

RNAi was performed by the feeding method, as described (Timmons et al., 2001). *E. coli* strain HT115 was transformed with either negative control vector pPD129.36 or plasmid targeting *vps-4* (Vidal library clone (Rual et al., 2004) - Y34D9A_152.a), *tsg-101* (Vidal library clone (Rual et al., 2004) - C09G12.9) or *hmr-1*, which results in dead embryos and was used as a positive control for RNAi efficacy (Chihara and Nance, 2012). L4 larvae (WT: FT1016, RAB-35^PGC-^: FT2561) were placed on RNAi plates with bacterial lawns and fed for 48 hours at 23°C. Gravid mothers were then dissected to release embryos in M9 buffer, which were allowed to hatch overnight before scoring for persistent lobes.

### Microscopy

Two-cell embryos released in M9 buffer from dissected adults were mounted on 4% agarose pads and incubated at 25°C until image acquisition began. Images were acquired using a Leica SP8 laser scanning confocal microscope with a 63X 1.4NA water objective and HyD detectors (standard counting mode). Images were acquired in 400nm slices at 400Hz and with a line accumulation of 2.

### Image acquisition intervals

**PIP_2_^END^/ PIP_2_^PGC^/ GFP11-RAB-35^SOMA^**: 10 minute, 38 second intervals. **GFP_11_-RAB-35^PGC^/TSG-101-GFP_11_**: 6 minute, 30 second intervals.

### Image analysis

**PIP_2_^END^/ PIP_2_^PGC^/ GFP_11_-RAB-35^SOMA^**: 2 µm lines were drawn such that +1 µm centered at the lobe membrane and extended into the endodermal cytoplasm (see Fig. 4A). **GFP11-RAB-35^SOMA^**: lines of interest were drawn to avoid puncta of GFP11-RAB-35^SOMA^ scattered through the endodermal cytoplasm and in proximity to the lobe; **PIP_2_^END^**: lines of interest were drawn to avoid endodermal cell membranes, as these membranes are additionally rich in PIP_2_^END^ signal. **TSG-101-GFP_11_**: ‘Intensity within lobes’ (Fig. 6D) was calculated by drawing a ROI around the periphery of the lobe and calculating the mean intensity. TSG-101-GFP_11_ puncta were analyzed by applying a ‘MaxEntropy’ (Fiji) threshold to images prior to the ‘3D Objects Counter’ (Fiji) function analysis (Fig. S5A-C).

**Image display:** Figures 1D’-D’’, 1E’-E’’, 2B-F’’, 3B-C’’, 4B-C’’, 4E-F’’, 5A-D’’, 6A-B’’: Micrographs were first rotated and cropped (Fiji) before image resolution (300ppi) was adjusted (Adobe Photoshop). Finally, brightness levels were adjusted (Fiji).

**Table S1.**
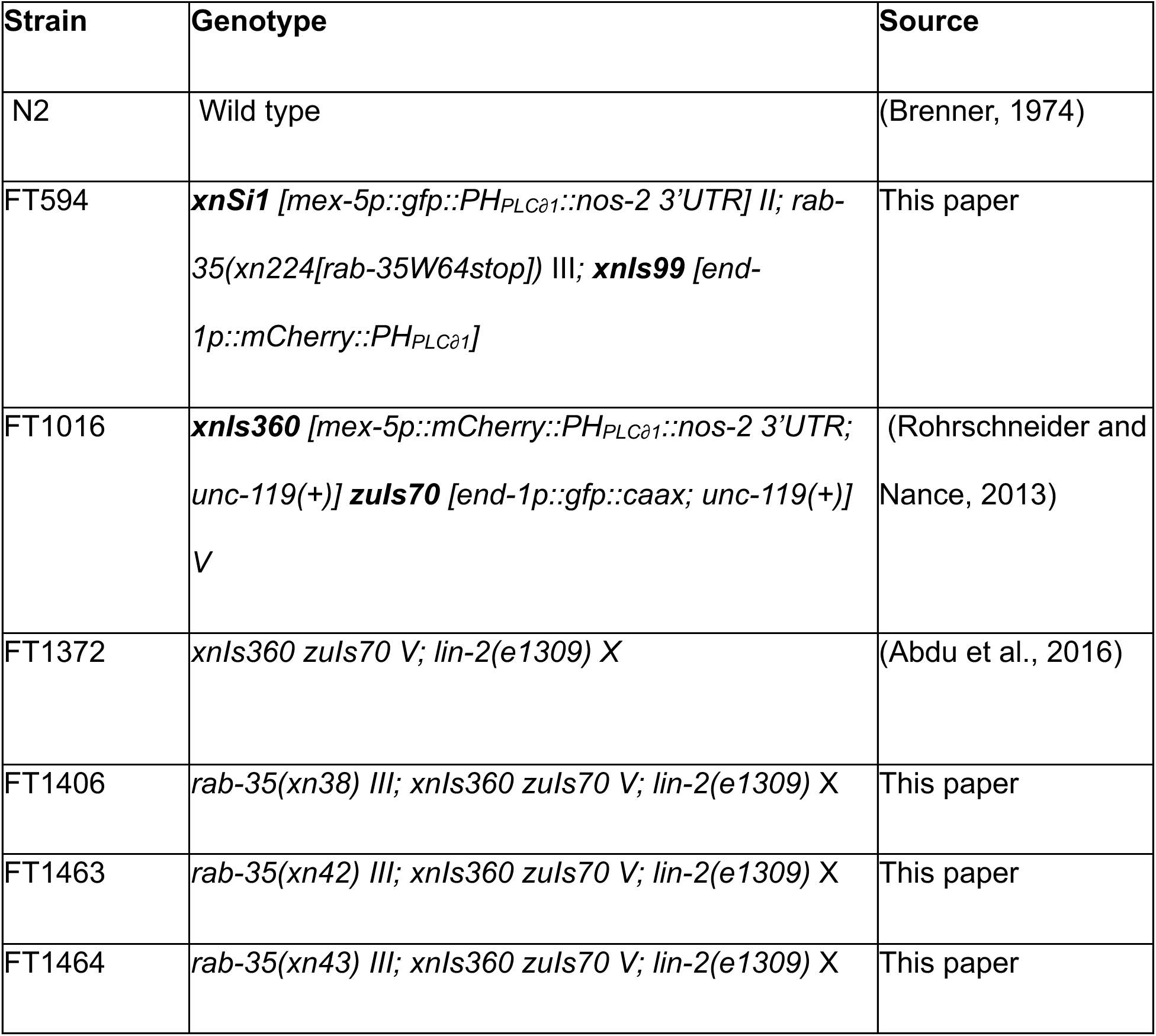
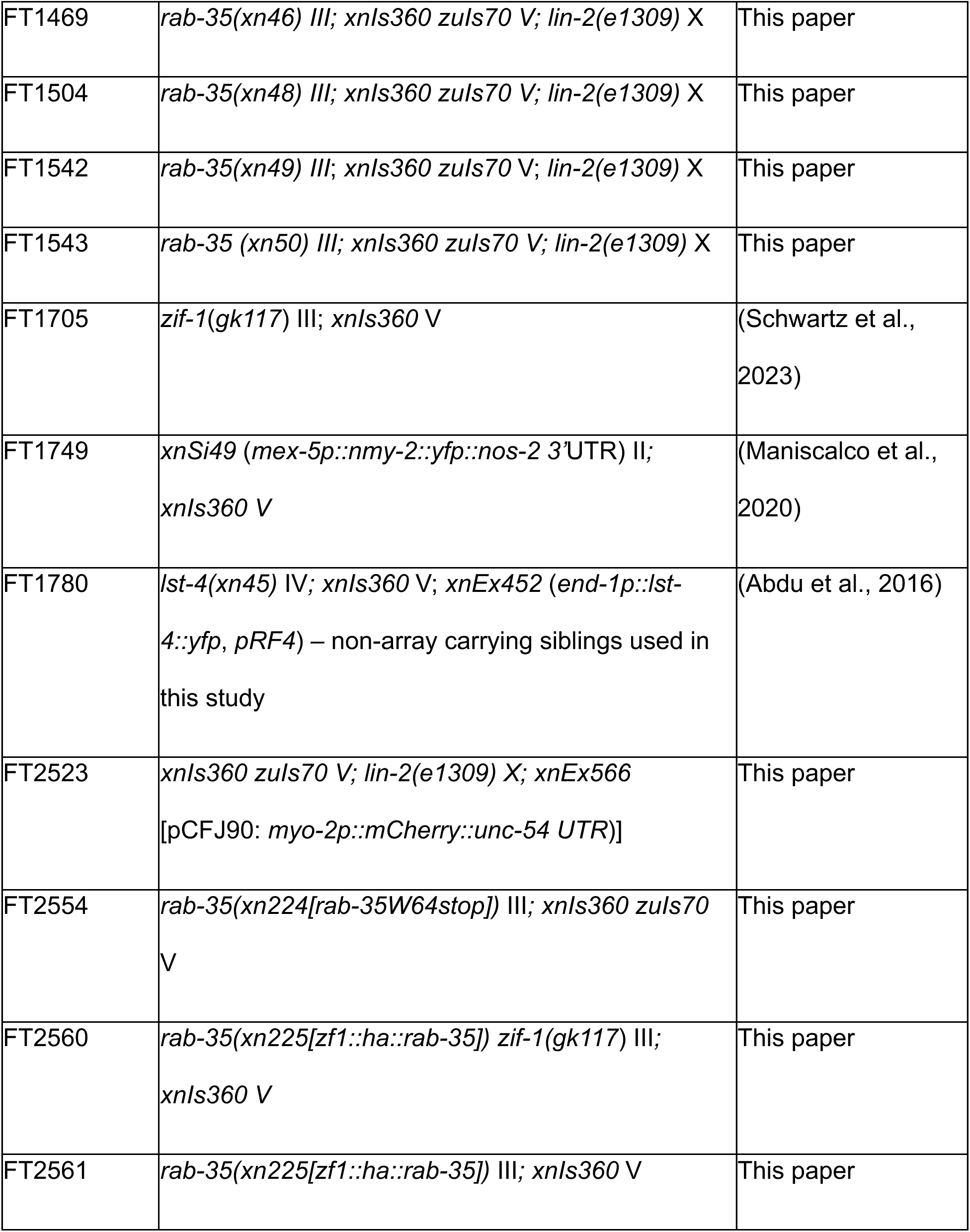
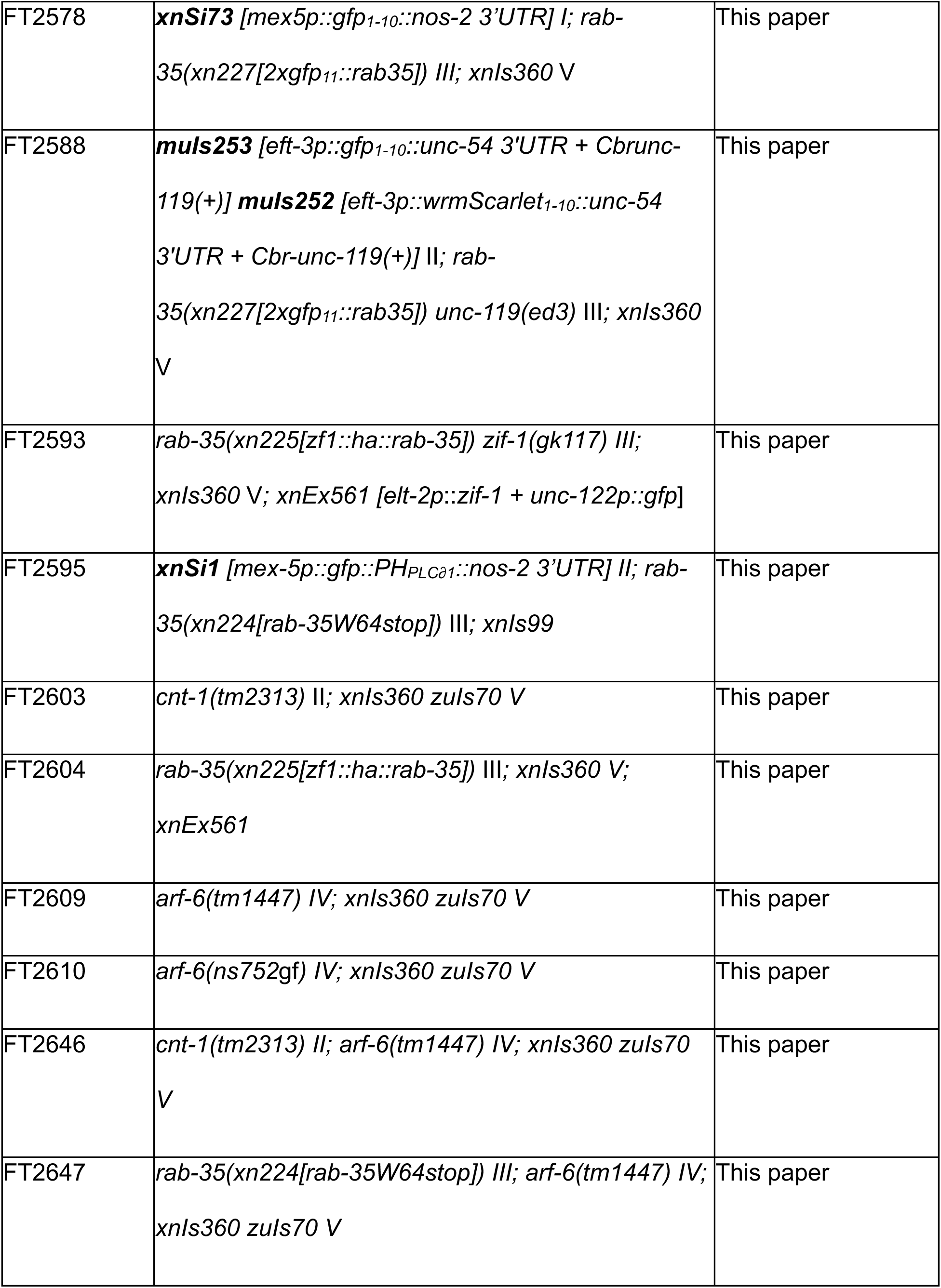
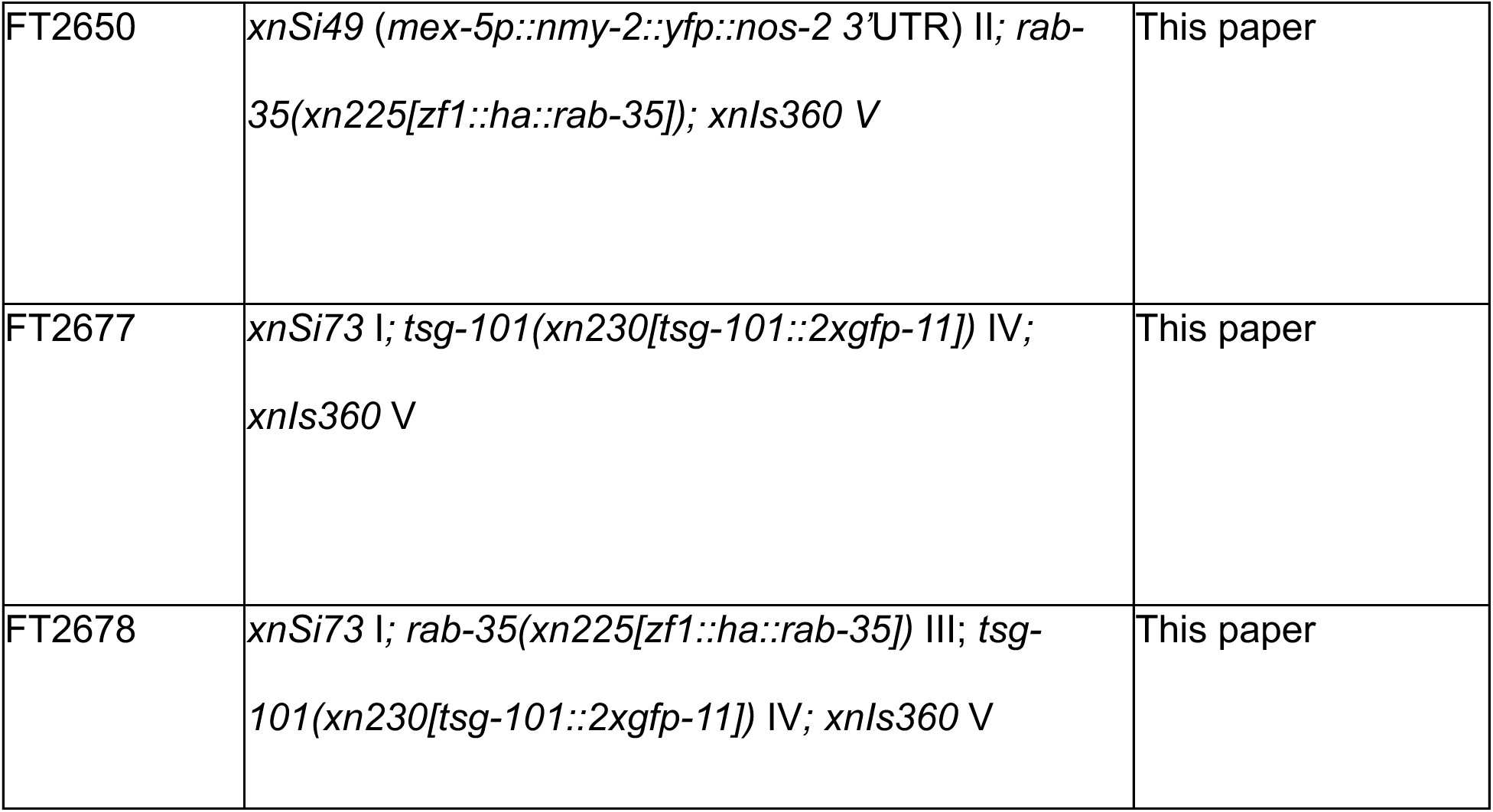
Worm Strains.

**Table S2.**
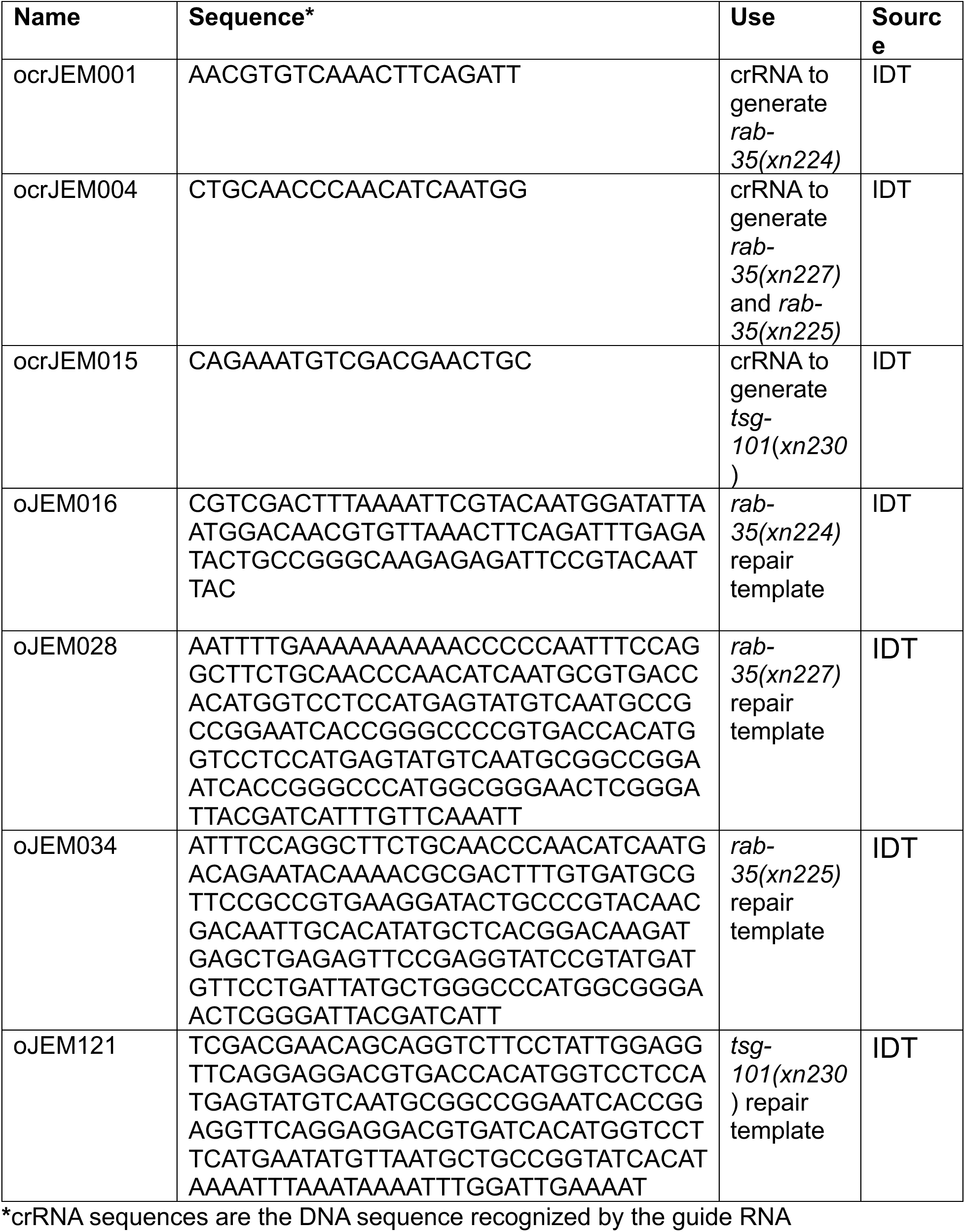
Sequence-based Reagents.

**Figure S1.**
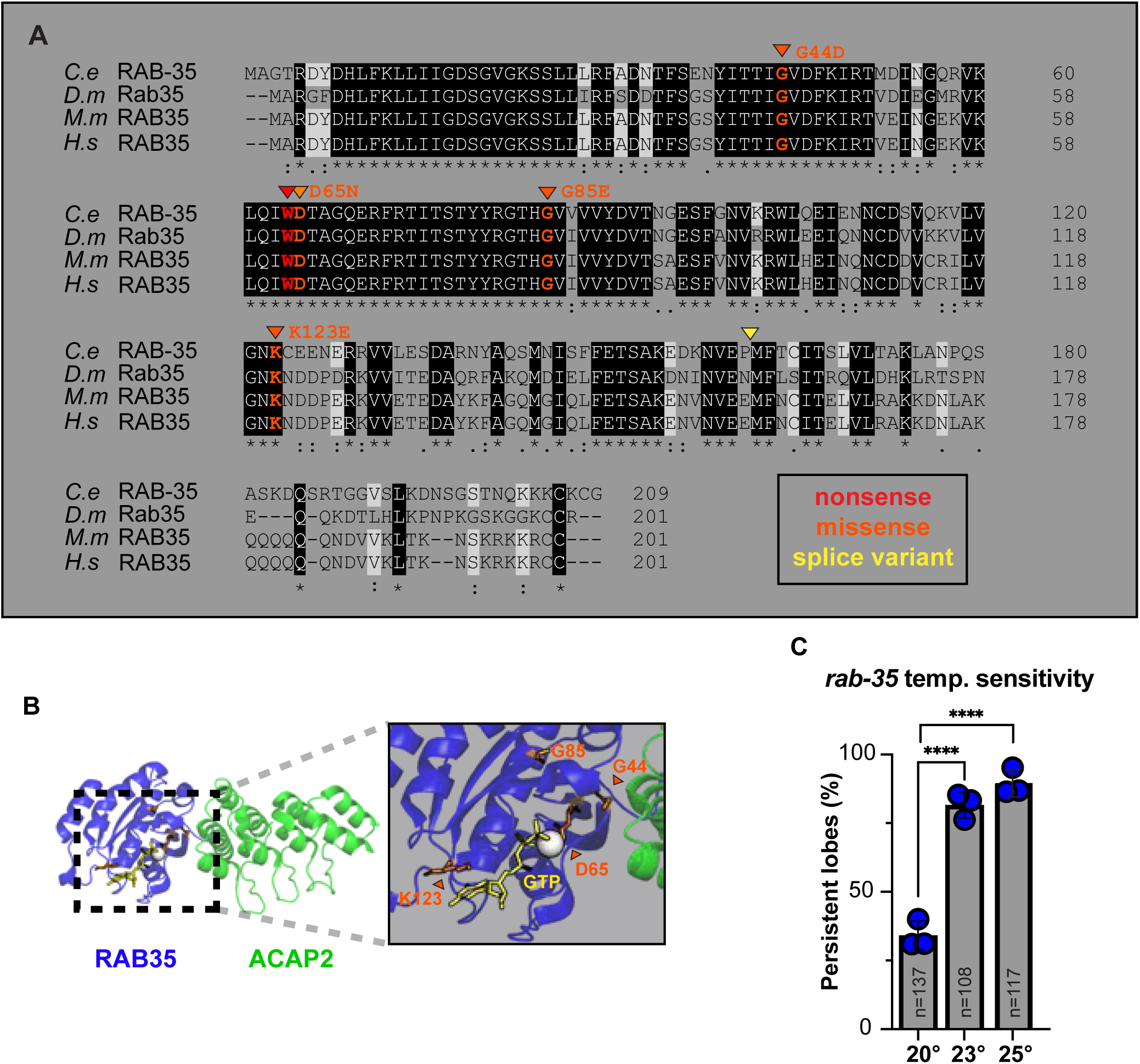
*rab-35* mutations and temperature sensitivity. **(A)** Protein sequences of RAB-35 homologues aligned using UniProt ‘align’ tool. Species abbreviations: *C.e* (*Caenorhabditis elegans*), *D.m* (*Drosophila melanogaster*), *M.m* (*Mus musculus)*, *H.s* (*Homo sapiens*). Shading/symbols: identical residues (black shading, asterisk below); strongly similar residues (period below); weakly similar residues (colon below); conserved residues in three of four species (white shading). Predicted changes introduced in *rab-35* mutant alleles are shown and colored as indicated in the legend. **(B)** PyMOL image of crystal structure of human RAB35/ACAP2 interaction (Protein Data Bank: 6IF3) (Lin et al., 2019). RAB35 (blue) and its effector ACAP2 (green). Inset: GTP-binding pocket and effector binding site. GTP (yellow), Mg^2+^ (white ball). The positions of *C. elegans* RAB-35 missense mutations are mapped onto the *H.s* RAB35 structure (orange); amino acid positions shown refer to *C. elegans* RAB-35. **(C)** Penetrance of the *rab-35* persistent lobe phenotype at the indicated temperature. Circles: percent of L1 with the phenotype in independent replicates (n≥34 animals each replicate); gray box, average of experimental replicates; error bars: SEM; n-values: total number of animals examined across all replicates. ****p-value ≤ 0.0001, Fisher’s exact test.

**Figure S2.**
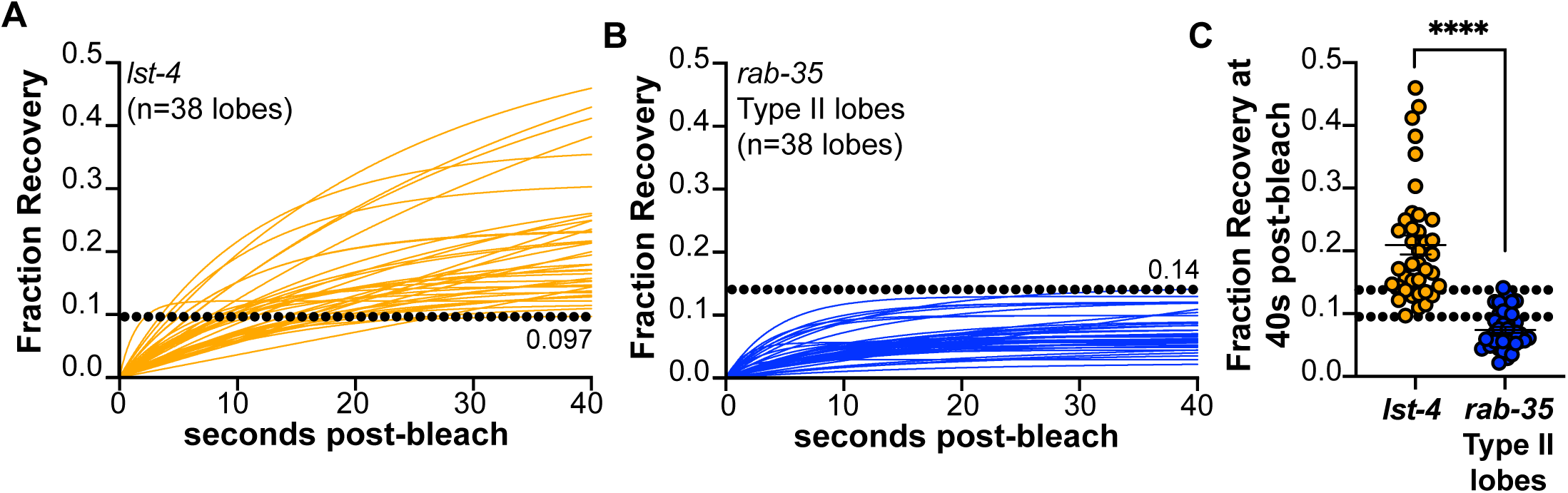
Calibration curves for FRAP assays. **(A-B)** Fluorescence recovery curves, fit to one-phase association models, of individual persistent lobes (see Methods). Lobes in (A) are from *lst-4* mutants, which block trogocytosis at the scission step. Lobes in (B) are an equal-size subset of Type II *rab-35* persistent lobes that were visually inspected to ensure that they were present well within endoderm and lacked a connection to the PGC cell body. Upper and lower range values indicated on graphs with dotted lines. **(C)** Fraction recovery values at 40 seconds post-bleach of curves shown in (A-B). Circles: individual persistent lobes; error bars: SEM; dotted lines: the lower (*lst-4*) and upper (*rab-35* Type II persistent lobes) limits of fraction recovery curves. ****p-value < 0.0001, unpaired nonparametric Mann-Whitney t-test.

**Figure S3.**
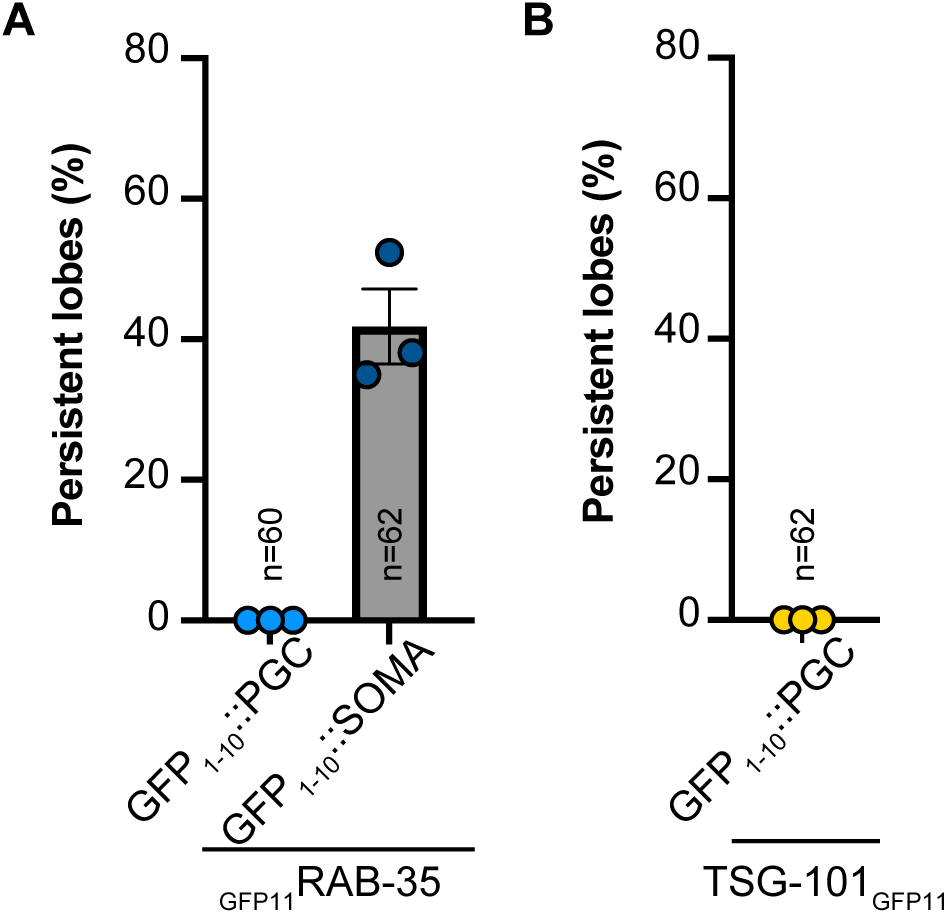
Testing functionality of GFP_11_-tagged *rab-35* and *tsg-101* alleles. **(A-B)** Penetrance of persistent lobe phenotype in L1 of the given genotype. Circles: percentage persistent lobes within a biological replicate (n ≥ 20 animals); error bars: SEM; n- value: number of larvae examined across all replicates.

**Figure S4.**
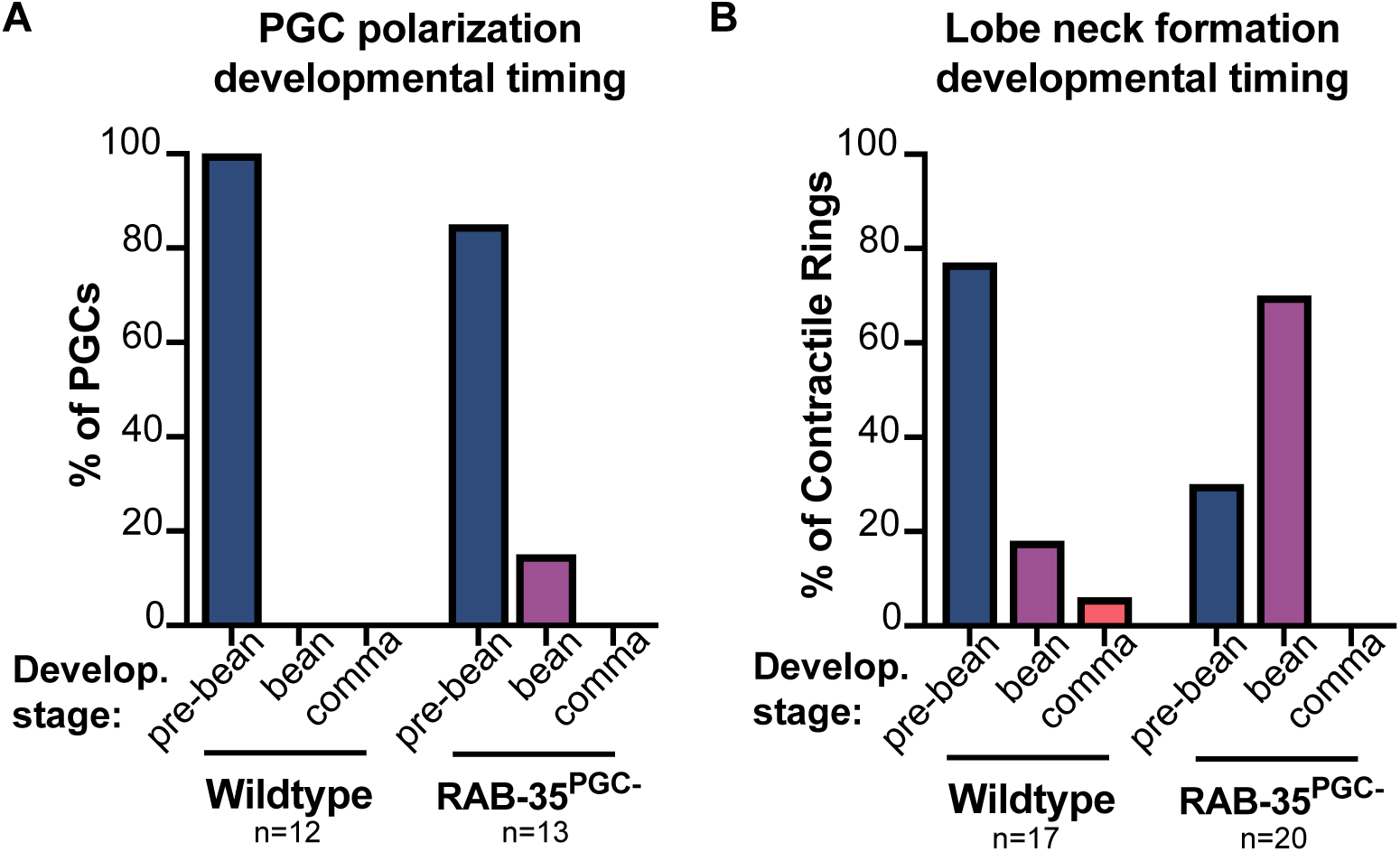
RAB-35 is not required for PGC lobe formation. **(A)** Developmental timing of PGC polarization. Bars: percent of PGCs that polarize at three different developmental stages in WT versus RAB-35^PGC-^ embryos. n-values: total number of PGCs analyzed across ≥ 2 biological representatives. **(B**) Developmental timing of lobe neck formation. Bars: percent of contractile rings that form at three different developmental stages in WT versus RAB-35^PGC-^ embryos. n-values: total number of PGCs analyzed across ≥ 2 biological representatives.

**Figure S5.**
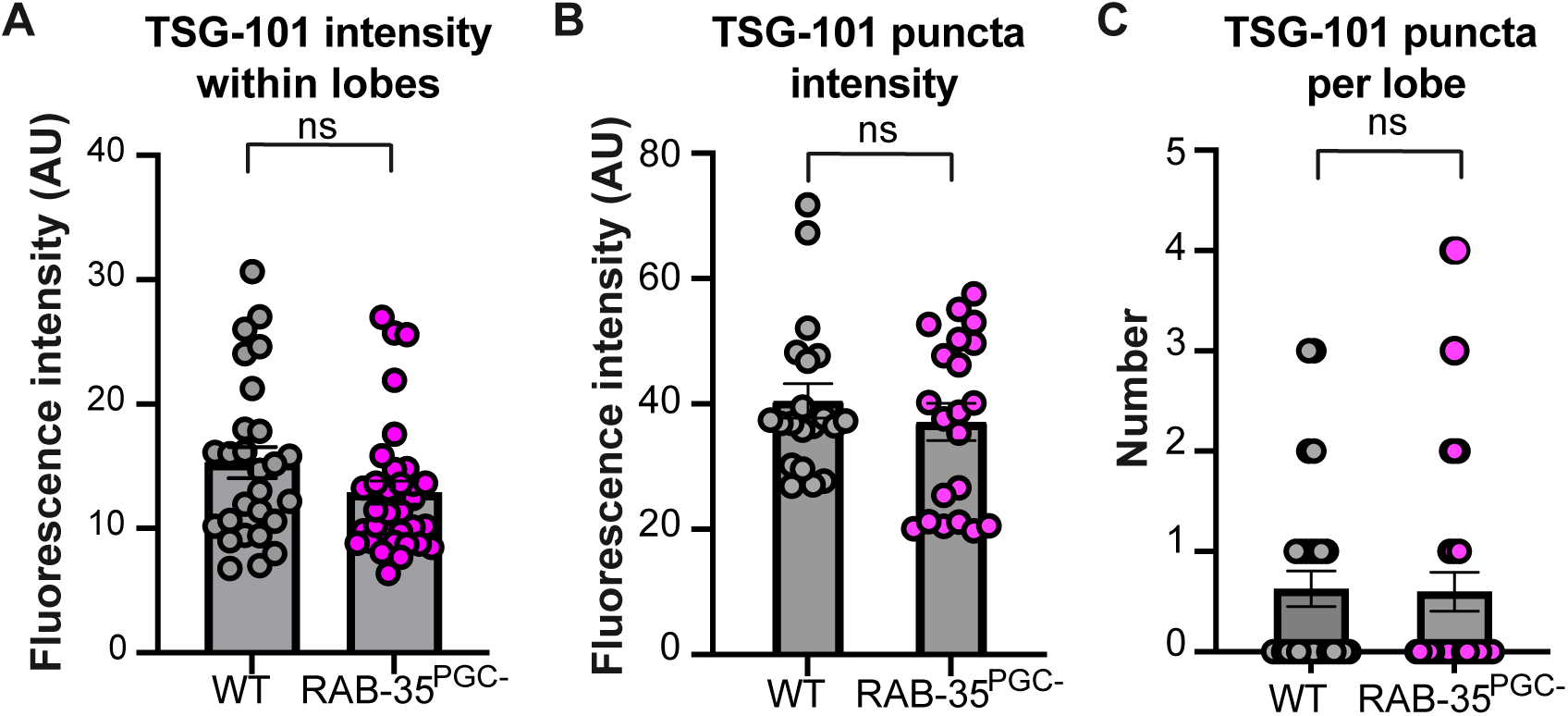
RAB-35 does not influence TSG-101^PGC^ expression or puncta formation. **(A)** TSG-101^PGC^ intensity within lobes. Circles, fluorescence intensity (AU) within a single lobe; bars, average fluorescence intensity of all lobes; error bars: SEM; n-values: number of lobes examined across ≥ 3 biological replicates. ns, p-value >0.05, two-tailed unpaired t-test. **(B)** TSG-101^PGC^ puncta intensity. Circles, fluorescence intensity (AU) of individual puncta; bars, average fluorescence intensity of all puncta; error bars: SEM; n-values: number of lobes examined across ≥ 3 biological replicates. ns, p-value >0.05, two-tailed unpaired t-test. **(C)** TSG-101 puncta per lobe. Circles, number of puncta identified within an individual lobe; bars; average number of puncta identified across lobes; error bars: SEM; n-values: number of lobes examined across ≥ 3 biological replicates. ns, p-value >0.05, two-tailed unpaired t-test.

